# Polysaccharide release factors drive lipid-carrier recycling in bacteria

**DOI:** 10.64898/2026.10.14.759067

**Authors:** Henri Voedts, Ombeline Verstraete, Rodolphe Auger, Delphine Patin, Jean Jacobs, Guido T. Bommer, Michel Arthur, Jean-Emmanuel Hugonnet, Bogdan I. Iorga, Thierry Touzé, Jean-François Collet

## Abstract

Gram-negative bacteria build their cell envelope by transporting various sugar-based precursors across the inner membrane on the shared lipid carrier undecaprenyl (also known as C55). Because the cellular pool of undecaprenyl is finite and used by multiple competing pathways, it must be efficiently regenerated to sustain envelope assembly. Yet the enzymes that release polysaccharide cargo from the lipid carrier have remained elusive, leaving a major gap in our understanding of bacterial envelope biogenesis. Here we identify three periplasmic enzymes in *Escherichia coli* — UprA, UprB, and UprC — that hydrolyze the bond linking polysaccharide precursors to undecaprenyl. UprA releases elongating peptidoglycan glycan strands from the inner membrane, catalyzing the long-sought terminal step of peptidoglycan synthesis. UprB cleaves the undecaprenyl-linked initiating unit common to the enterobacterial common antigen and O-antigen pathways, while UprC is the likely release factor for the colanic acid pathway. These enzymes define a widespread Upr-like protein family (formerly DUF218) as polysaccharide release factors, revealing a unifying mechanism by which bacteria coordinate envelope assembly. Because all three Upr-like proteins share a conserved catalytic core, this family represents a potential target for antibiotics that simultaneously disrupt multiple envelope pathways.

## INTRODUCTION

The cell envelope of Gram-negative bacteria is a three-layered structure essential for protecting the cell from internal turgor pressure and environmental stress ^1,2^. It consists of an inner membrane and an outer membrane, which delimit the periplasmic space containing the peptidoglycan layer. Peptidoglycan is a net-like macromolecule that surrounds the cell, provides mechanical strength, and enables the buildup of a counterbalancing periplasmic turgor pressure that stabilizes the inner membrane^3^.

Cell envelope biogenesis relies on pathways that move building blocks synthesized in the cytoplasm across the inner membrane and, for those contributing to peptidoglycan and outer membrane biogenesis, extract them from the inner membrane, and guide them to their final destination^4^. Within this framework, enabling hydrophilic intermediates to traverse the inner membrane and engage in assembly reactions on the periplasmic side is a major challenge. A widely conserved solution is the use of lipid-linked precursors, with the undecaprenyl lipid carrier (hereafter C55; phosphorylation states specified as C55-P or C55-PP when relevant) serving as a central scaffold (**Fig. 1a**). C55 is an essential long-chain isoprenoid enabling the transport of multiple envelope glycans. Substrates are attached to C55-P via a diphosphate linkage, generating C55-PP-linked intermediates that are flipped across the inner membrane and serve as platforms for periplasmic polymerization^4^. After cargo transfer, the lipid carrier must be regenerated from its diphosphate form to C55-P^5^ and returned to the cytoplasmic face of the inner membrane to restart the cycle. Recent studies identified two broadly conserved protein families — DedA and DUF368-containing proteins — as candidate C55-P translocases that mediate this last step^6,7^, thereby filling a critical gap in our understanding of lipid-carrier recycling.

**Figure 1.**
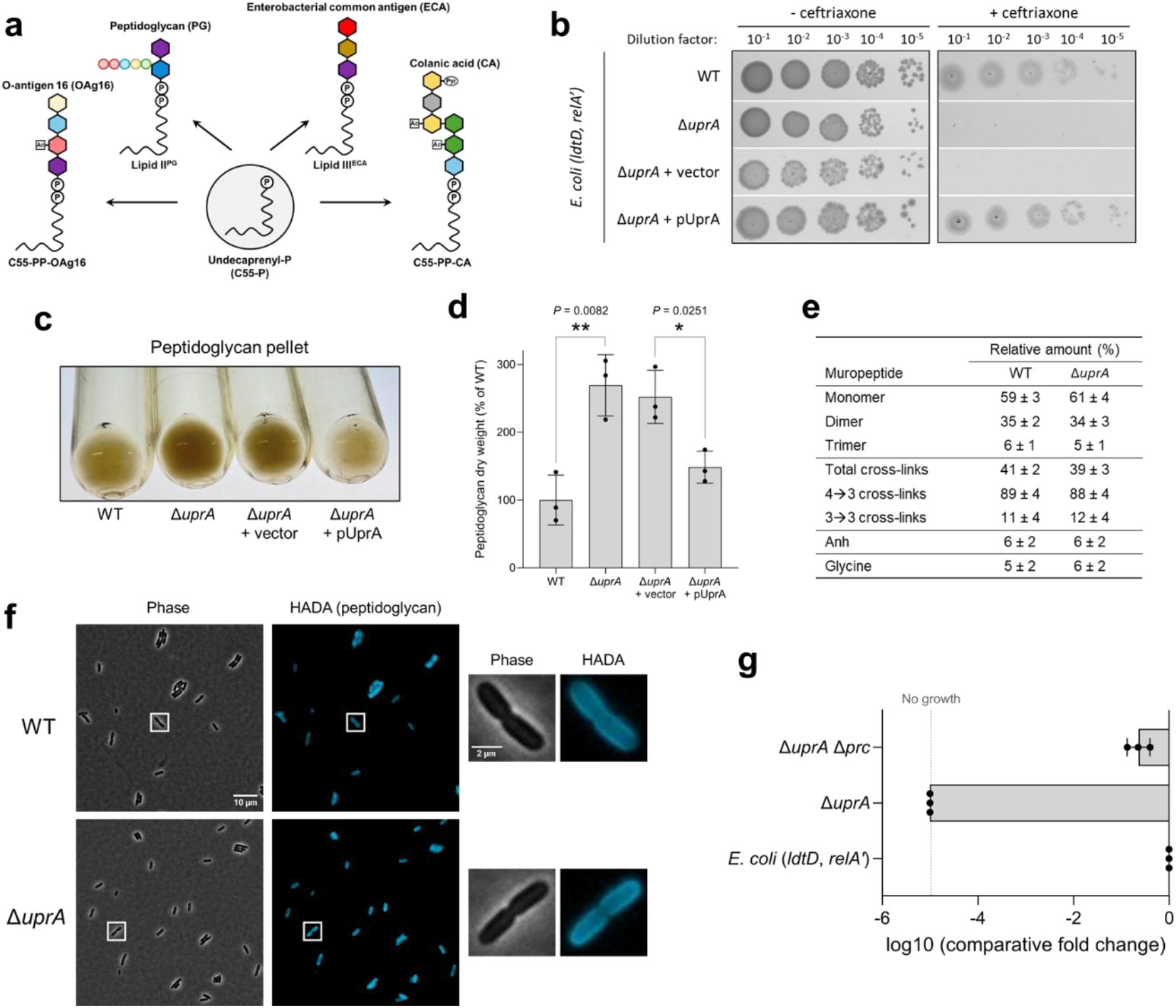
UprA is linked to peptidoglycan metabolism. (**a**) Schematic illustrating the envelope polysaccharide synthesis pathways relying on C55-P. (**b**) Plating efficiency of *E. coli* (*ldtD*, *relA’*) and its derivatives harboring the deletion of *uprA* and plasmids for complementation. Growth was tested in the presence of ceftriaxone at 2 µg/mL (+ ceftriaxone) or in the absence of the drug (-ceftriaxone). BHI agar plates contained 40 µM IPTG and 1% L-arabinose for induction of *ldtD* and *relA’*, respectively. (**c**) Picture of the peptidoglycan pellets of *E. coli* BW25113 (WT) and derivatives after extraction. (**d**) Dry weight quantification of peptidoglycan pellets of *E. coli* BW25113 (WT) and derivatives. The vertical bars correspond to the mean of the peptidoglycan dry weight relative to WT, the error bars represent the standard deviation, and the dots represent individual data (n = 3 biological replicates). (**e**) Comparative table of peptidoglycan composition of *E. coli* BW25113 (WT) and the derivative harboring the deletion of *uprA*. Data represent the mean ± standard deviation (n = 3 biological replicates). Anh, anhydromuropeptide. (**f**) Microscopic analysis of *E. coli* BW25113 (WT) and its derivative harboring the deletion of *uprA*. White squares indicate the zoomed cell on the right side of the panel. (**g**) Plating efficiency of *E. coli* (*ldtD*, *relA’*) and its derivatives harboring the deletion of *uprA* and *prc.* Growth efficiencies were determined relative to the growth of the parental strain *E. coli* (*ldtD*, *relA’*). Growth was tested in the presence of 2 µg/mL ceftriaxone. BHI agar plates contained 40 µM IPTG and 1% L-arabinose for induction of *ldtD* and *relA’*, respectively. The horizontal bars correspond to the mean, the error bars represent the standard deviation, and the dots represent individual data (n = 3 biological replicates).

A key C55-dependent precursor is lipid II^PG^ (**Fig. 1a**), which fuels peptidoglycan synthesis. At the cytoplasmic side of the membrane, the UDP-linked *N*-acetylmuramic acid (MurNAc) pentapeptide is attached to C55-P to form lipid I^PG^, then converted to lipid II^PG^ by addition of *N*-acetylglucosamine (GlcNAc; **Extended Data Fig. 1a**). Lipid II^PG^ is subsequently flipped to the periplasmic side of the inner membrane, where glycosyltransferases polymerize the disaccharide-peptide unit into glycan strands that are further cross-linked by transpeptidases^8^ (**Extended Data Fig. 1a**). Critically, peptidoglycan is not the only pathway that depends on C55. In *Escherichia coli*, C55 also supports the synthesis of the enterobacterial common antigen (ECA), a conserved carbohydrate found in the envelope^9^ (**Extended Data Fig. 1b**); the O-antigen, assembled on C55 before ligation to LPS and displayed at the cell surface^10^ (**Extended Data Fig. 1c**); and colanic acid, a heteropolysaccharide that can be conjugated to LPS (M-antigen) or secreted extracellularly^11^ (**Extended Data Fig. 1d**). Thus, multiple essential pathways draw on a single lipid carrier.

Despite its central role in envelope biogenesis, the cellular pool of C55 is intrinsically limited (about 10^5^ molecules per *E. coli* cell)^12,13^. Its availability therefore constitutes a critical point of coordination between envelope pathways: accumulation of lipid-linked intermediates in one pathway can sequester the carrier and indirectly starve the others^14^. Yet, how bacteria regulate the distribution and recycling of this finite resource has remained largely undefined. Although the translocases that return unloaded C55-P to the cytoplasmic face of the inner membrane were recently identified^6,7^, the enzymatic mechanisms that disengage polysaccharide intermediates from the lipid carrier —thereby regenerating C55 for reuse— remain unknown. This unresolved step not only constrains steady-state envelope synthesis, but becomes particularly critical when lipid-linked intermediates are incomplete, stalled, or fail to progress through downstream steps. Nowhere is the need for such a release reaction more evident than in peptidoglycan synthesis: glycosyltransferases elongate glycan strands by transferring the nascent chain onto newly translocated lipid II^PG^ molecules, a process that leaves the growing polymer covalently tethered to C55 (**Extended Data Fig. 1a**). How these membrane-anchored chains are released into the sacculus has remained a mystery, representing one of the last unresolved steps in peptidoglycan assembly. In the other C55-dependent pathways, some downstream transfer reactions have been characterized, including ligation to LPS (**Extended Data Fig. 1b-d**). However, important steps remain unclear, notably how free colanic acid is released and how ECA is attached to phosphatidylglycerol (**Extended Data Fig. 1b,d**). More generally, how cells clear aberrant or stalled lipid-linked intermediates and thereby recycle the carrier remains unresolved. Thus, despite decades of work on cell-envelope biogenesis, the enzymes that actively liberate C55-linked polymers to prevent carrier sequestration have remained elusive.

Here, we define the biological function of three periplasmic *E. coli* enzymes — UprA (formerly SanA), UprB (formerly ElyC), and UprC (formerly YgjQ) — as hydrolases of C55-PP-linked polysaccharide precursors. UprA hydrolyzes lipid II^PG^ and releases elongating glycan strands from the inner membrane, catalyzing the long-sought terminal step of peptidoglycan synthesis. UprB cleaves the C55-PP-GlcNAc linkage shared by the ECA and O-antigen biosynthetic routes and, accordingly, relieves C55 sequestration when O-antigen precursors accumulate in the periplasm. UprC, in turn, limits the buildup of intermediates in the colanic acid pathway, identifying it as the likely release factor for this third C55-dependent pathway. These proteins belong to the widespread DUF218 family, which we redesignate as the Upr-like family (for <u>u</u>ndecaprenyl-diphosphate-<u>p</u>olysaccharide <u>r</u>elease factor). Genetic and biochemical evidence reveals that Upr-like enzymes are partitioned across envelope pathways *in vivo*, despite displaying partially overlapping substrate scope *in vitro*. Together, these findings uncover a unifying enzymatic principle for polysaccharide release and lipid-carrier recycling across bacterial envelope biogenesis.

## RESULTS

### UprA restrains peptidoglycan accumulation and acts on lipid-linked precursors

We previously identified *uprA* (formerly *sanA*) as being required for LdtD-mediated β-lactam resistance^15^, linking this gene of unknown function to peptidoglycan metabolism. LdtD is an L,D-transpeptidase that cross-links peptidoglycan peptide stems through 3→3 linkages^16^. When the D,D-transpeptidases responsible for canonical 4→3 cross-links are inhibited by β-lactams such as ceftriaxone, LdtD can sustain growth by functionally replacing them — but only in the presence of *uprA*^15,17^ (**Fig. 1b**). Beyond this genetic association, UprA — an inner membrane protein with a large periplasmic domain (**Extended Data Fig. 2**) — has been implicated in envelope integrity^18^, yet its molecular function has remained elusive for three decades. A critical clue came from the protein family to which UprA belongs. The Upr-like family also includes UprB (formerly ElyC; **Extended Data Fig. 2**), a periplasmic protein directly implicated in C55 homeostasis: loss of UprB leads to a marked increase in ECA production with a concomitant decrease in peptidoglycan synthesis, a phenotype attributed to sequestration of the shared C55 lipid carrier by ECA precursors^19,20^. The direct involvement of UprB in C55-dependent ECA homeostasis, together with the genetic link between UprA and peptidoglycan — another pathway critically dependent on C55 — raised a clear possibility: Upr-like proteins might act directly on C55-linked polysaccharide intermediates. We therefore set out to test whether members of this family function as release factors that liberate envelope polymers from the lipid carrier.

To explore this hypothesis, we first examined the consequences of *uprA* deletion on peptidoglycan architecture and composition. Quantification revealed an approximately 2.5-fold increase in peptidoglycan in Δ*uprA* cells relative to wild type (**Fig. 1c,d**), consistent with a recent report of increased radioactive *meso*-diaminopimelic acid incorporation upon *uprA* deletion^21^. Despite this substantial increase, muropeptide analysis by liquid chromatography-mass spectrometry (LC-MS) showed no major changes in peptidoglycan fragment composition (**Fig. 1e; Extended Data Fig. 3**), and labelling with fluorescent D-amino acids^22^ uncovered no detectable morphological abnormalities (**Fig. 1f**). Together, these observations indicate that loss of UprA causes peptidoglycan to accumulate without altering its structural organization, consistent with a role in restraining or terminating synthesis rather than in assembly or remodeling. Independent support for this idea was provided by a suppressor analysis; spontaneous mutations that restored LdtD-mediated β-lactam resistance in Δ*uprA* cells mapped to *prc* (**Fig. 1g**), encoding a periplasmic protease that normally degrades peptidoglycan hydrolases^23–25^. Null mutations in *prc* have been shown to stabilize these hydrolases, increasing peptidoglycan turnover and thereby compensating for the excess accumulation.

We next asked whether UprA acts directly on lipid-linked peptidoglycan precursors. Lipid II^PG^, either purified from *E. coli* or synthesized *in vitro*, was incubated with purified UprA and reaction products monitored by LC-MS. UprA efficiently hydrolyzed both substrates, releasing GlcNAc-MurNAc-pentapeptide from C55-PP (**Fig. 2a,b**). The reaction proceeded in a time- and concentration-dependent manner (**Fig. 2c**). Thus, UprA is a lipid II^PG^ hydrolase. Beyond this canonical substrate, UprA also hydrolyzed a minor lipid II^PG^ species that contains a tetrapeptide-stem that was present in the synthetic preparation (**Fig. 2d**), indicating tolerance to variation in the peptide stem. UprA even hydrolyzed lipid I^PG^ *in vitro* (**Fig. 2e**), demonstrating that it can liberate C55-PP from incomplete peptidoglycan intermediates. Together, these activities suggest a quality-control role, in which UprA rescues C55 from immature or abortive precursors when polymerization is delayed or fails.

**Figure 2.**
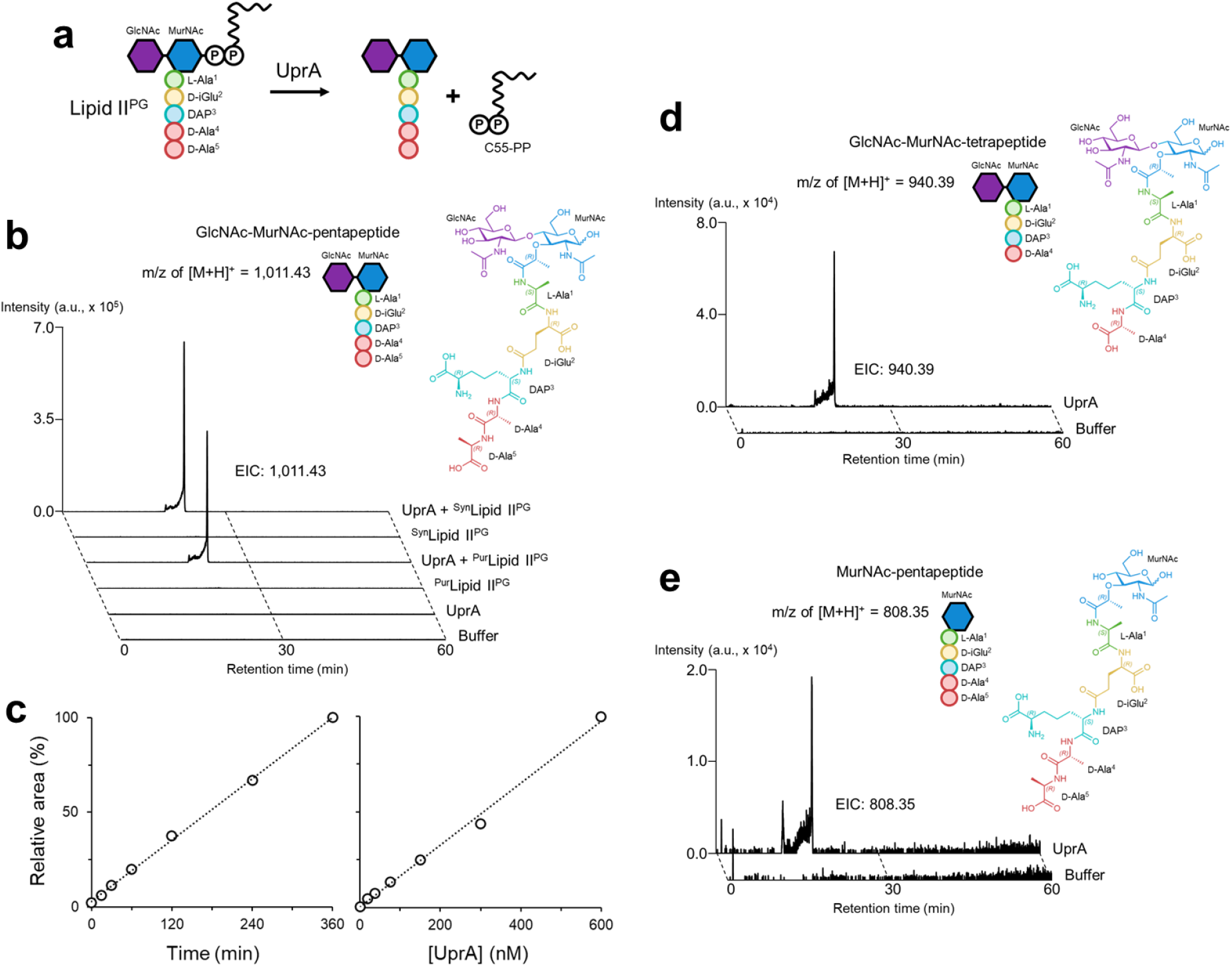
UprA catalyzes the release of the peptidoglycan subunits from the C55-PP lipid carrier. (**a**) Schematic illustrating the reaction catalyzed by UprA. (**b**) Extracted ion chromatograms (EICs) of the *in vitro* enzymatic reaction of UprA with purified lipid II^PG^ (^Pur^Lipid II^PG^) and synthesized lipid II^PG^ (^Syn^Lipid II^PG^). The structure of the detected product GlcNAc-MurNAc-pentapeptide is shown for reference. a.u., arbitrary unit. (**c**) The *in vitro* enzymatic activity of UprA is time- and concentration-dependent. The relative area under the curve of GlcNAc-MurNAc-pentapeptide was obtained by comparing the area to the maximum (*i.e.* 360 min or 600 nM). (**d**) Extracted ion chromatograms (EICs) of the *in vitro* enzymatic reaction of UprA with synthesized lipid II^PG^ partially processed by a contaminant D,D-carboxypeptidase. The structure of the detected product GlcNAc-MurNAc-tetrapeptide is shown for reference. a.u., arbitrary unit. (**e**) Extracted ion chromatograms (EICs) of the *in vitro* enzymatic reaction of UprA with synthesized lipid I^PG^. UprA was directly incubated with C55-P, UDP-MurNAc-pentapeptide, and MraY. The structure of the detected product MurNAc-pentapeptide is shown for reference. a.u., arbitrary unit.

Protenix^26^ structural predictions revealed a well-defined cavity within the Upr-like domain of UprA (**Fig. 3a**). Structural modeling positioned lipid II^PG^ within this cavity, orienting the scissile phosphoacetal bond near a cluster of conserved residues — T^125^, H^149^, R^152^, and E^182^ (**Fig. 3a,b; Extended Data Fig. 2**). To test the catalytic role of these residues, we monitored UprA activity using a fluorescence-based assay^27,28^: dansylated-lipid II^PG^ fluoresces in the hydrophobic environment promoted by the C55 lipid tail, whereas hydrolysis releases the GlcNAc-MurNAc-pentapeptide carrying the dansyl moiety into the aqueous phase, quenching the signal. Alanine substitutions at all four positions abolished UprA activity in this assay (**Fig. 3c**) and failed to restore β-lactam resistance in Δ*uprA* cells (**Fig. 3d**). By contrast, conservative substitutions — T^125^S and E^182^D — restored β-lactam resistance (**Fig. 3d**), consistent with a direct catalytic role for the corresponding residues. To define the catalytic mechanism, we performed the reaction in [^18^O]H_2_O and analysed the products by LC-MS. ^18^O incorporated specifically on the GlcNAc-MurNAc-pentapeptide product (**Fig. 3e**), establishing that water attacks the sugar side of the phosphoacetal bond rather than on the phosphate side. UprA activity required divalent Mg^2+^ or Ca^2+^ cations (**Fig. 3f**), similar to peptidoglycan glycosyltransferases for which divalent cations are required to stabilize the negative charges of the C55-PP moiety^29,30^. Vancomycin, which binds the D-Ala-D-Ala terminus of the lipid II^PG^ peptide stem, strongly inhibited hydrolysis (**Fig. 3g**). Modeling vancomycin into the active site explained this inhibition: vancomycin sterically excludes lipid II^PG^ from the catalytic cavity of UprA, displacing the scissile phosphoacetal bond from 4.8 Å to 11.7 Å from T^125^ (**Extended Data Fig. 4a-d**). This supports a model in which the lipid II^PG^ peptide stem must engage the catalytic cavity for hydrolysis.

**Figure 3.**
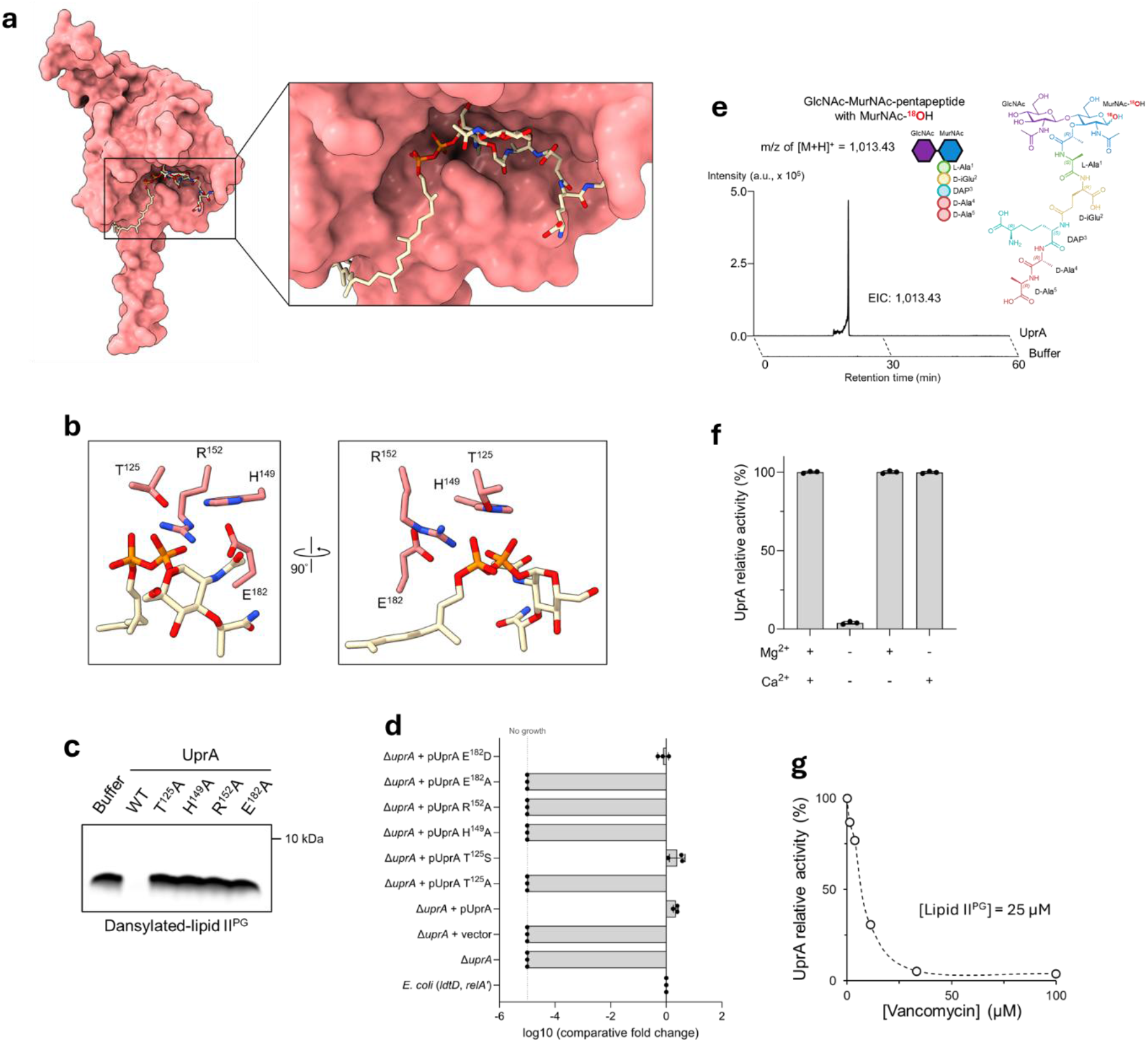
Biochemical characterization of UprA-mediated lipid II^PG^ hydrolysis. (**a**) Surface representation of the modeling of UprA with lipid II^PG^. The modeling shows entry of PP-GlcNAc-MurNAc-pentapeptide in UprA’s cavity containing the conserved residues of Upr-like proteins. (**b**) Stick representation on the conserved residues of UprA modelled with lipid II^PG^. For the sake of simplicity, only the C10-PP-MurNAc part of lipid II^PG^ is shown. (**c**) Fluorescence of synthesized dansylated-lipid II^PG^ incubated with UprA or catalytic-inactive variants. The dansyl moiety, attached to the third amino acid of the peptide stem, fluoresces in a hydrophobic environment promoted by the C55 lipid tail. A decrease in fluorescence indicates release of the dansyl moiety from the hydrophobic environment, consistent with cleavage of the glycan strands from the C55-PP. (**d**) Plating efficiency of *E. coli* (*ldtD*, *relA’*) and its derivatives harboring the deletion of *uprA* and plasmids for complementation with catalytic-inactive variants. Growth efficiencies were determined relative to the growth of the parental strain *E. coli* (*ldtD*, *relA’*). Growth was tested in the presence of 2 µg/mL ceftriaxone, and 50 µM DAPG for induction of *uprA* variants. BHI agar plates contained 40 µM IPTG and 1% L-arabinose for induction of *ldtD* and *relA’*, respectively. The horizontal bars correspond to the mean, the error bars represent the standard deviation, and the dots represent individual data (n = 3 biological replicates). (**e**) UprA catalyzes a hydrolysis reaction. Extracted ion chromatograms (EICs) of the *in vitro* hydrolysis of lipid II^PG^ by UprA in an aqueous buffer containing heavy-oxygen isotope [^18^O]H_2_O. The structure of the detected product GlcNAc-MurNAc-pentapeptide labelled with one atom of ^18^O is shown for reference. a.u., arbitrary unit. (**f**) The *in vitro* enzymatic activity of UprA requires Mg^2+^ or Ca^2+^. The horizontal bars correspond to the mean of the relative area of GlcNAc-MurNAc-pentapeptide, the error bars represent the standard deviation, and the dots represent individual data (n = 3 replicates). (**g**) UprA catalytic activity is prevented by the addition of vancomycin. The relative area of GlcNAc-MurNAc-pentapeptide was obtained by comparing areas to the area without vancomycin.

We next asked whether UprA also acts on mature peptidoglycan. UprA showed no detectable activity toward purified sacculi or soluble muropeptides, and direct assays demonstrated that it is neither a muramidase, a glucosaminidase, a lytic transglycosylase, a carboxypeptidase, nor an endopeptidase, ruling out canonical peptidoglycan hydrolase activities (**Extended Data Fig. 5**). Together, these results establish that UprA targets lipid-linked intermediates through hydrolysis of the phosphoacetal bond. Given that elongating glycan strands remain covalently tethered to C55^31,32^, this activity suggested a role in releasing nascent glycan chains from the membrane — the long-sought final step in peptidoglycan synthesis.

### UprA releases elongating glycan strands

To directly test this, we extended the dansylated-substrate approach to polymerized glycan strands. Lipid-linked strands were generated *in vitro* using PBP1bγ S^510^A, a variant that retains glycosyltransferase but not transpeptidase activity^33,34^, thereby producing uncross-linked products that remain anchored to C55 and fluoresce thanks to the hydrophobic environment. Addition of UprA caused a complete loss of fluorescence (**Fig. 4a**), indicating release of the strands into the aqueous phase. The same experiment conducted with the catalytic-inactive UprA T^125^A also showed a decrease in fluorescence, albeit to notably lesser extent. This residual activity persisted despite multiple rounds of purification and progressively shortened the glycan strands over time, suggesting the presence of a tightly co-purifying lytic transglycosylase contaminant active on uncross-linked glycan strands. Consistent with this activity on extended substrates, modeling of UprA in complex with a lipid-linked three-subunit glycan strand positioned the diphosphate and the first MurNAc within the catalytic cavity, with two distinct channels accommodating the peptide stem of the first subunit and the polymerized glycan chain, respectively (**Fig. 4b**). This two-channel architecture provides a structural rationale for how the same catalytic cavity can engage both lipid II^PG^ and an elongating glycan strand.

**Figure 4.**
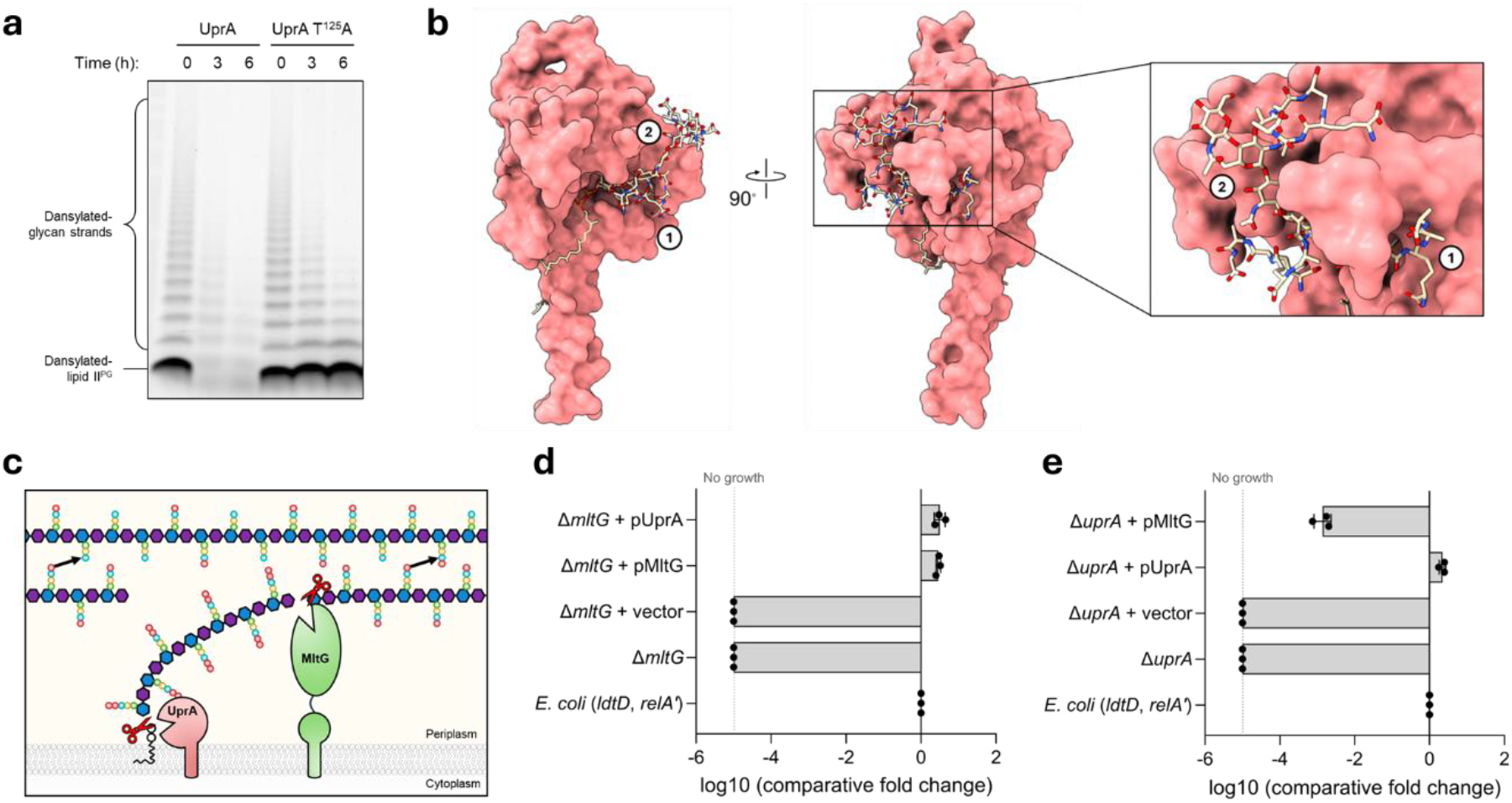
UprA catalyzes the last step of peptidoglycan synthesis: the release of the elongating glycan strand from the C55-PP lipid carrier. **(a)** Fluorescence of synthesized dansylated-lipid II^PG^ polymerized into glycan strands by PBP1bγ S^510^A and incubated with UprA or catalytic-inactive UprA T^125^A variant. The dansyl moiety, attached to the third amino acid of the peptide stem, fluoresces in a hydrophobic environment promoted by the C55 lipid tail. A decrease in fluorescence indicates release of the dansyl moiety from the hydrophobic environment, consistent with cleavage of the glycan strands from the C55-PP. **(b)** Surface representation of the modeling of UprA with a glycan strand resulting from the polymerization of three lipid II^PG^ molecules. The modeling shows entry of the diphosphate and the first subunit of the glycan strand in UprA’s cavity containing the conserved catalytic residues. Two channels accommodate (1) the peptide stem of the first subunit and (2) the polymerized glycan strand. **(c)** Schematic illustrating the reactions catalyzed by UprA and MltG. During the elongation process, the newly synthesized glycan strand is attached to the cytoplasmic membrane through the C55-PP lipid carrier. UprA releases the new glycan strand through hydrolysis of the bond between the C55-PP and the first MurNAc of the glycan strand. MltG release the new glycan strand through its lytic transglycosylase activity after the seventh subunit of the new glycan strand, leaving an engaged seven-subunits-long glycan strand linked to C55-PP^36^. **(d, e)** Plating efficiency of *E. coli* (*ldtD*, *relA’*) and its derivatives harboring the deletion of *mltG* (d) or *uprA* (**e**) and plasmids for complementation with *mltG* or *uprA*. Growth efficiencies were determined relative to the growth of the parental strain *E. coli* (*ldtD*, *relA’*). Growth was tested in the presence of 2 µg/mL ceftriaxone, and 50 µM DAPG for induction of *mltG* or *uprA*. BHI agar plates contained 40 µM IPTG and 1% L-arabinose for induction of *ldtD* and *relA’*, respectively. The horizontal bars correspond to the mean, the error bars represent the standard deviation, and the dots represent individual data (n = 3 biological replicates).

*In vivo*, the only enzyme previously implicated in glycan-strand release is the lytic transglycosylase MltG^35–37^, which is also required for LdtD-mediated β-lactam resistance^15^. MltG cleaves the glycan backbone of nascent peptidoglycan after the seventh subunit (**Fig. 4c**), distal to the C55 lipid carrier, releasing the newly synthesized glycan chain but leaving a committed elongation intermediate still attached to C55^36^ (**Fig. 4c**). UprA acts through a fundamentally different mechanism, hydrolyzing the lipid-glycan linkage itself to fully liberate the polymer from the membrane and the C55 lipid carrier (**Fig. 4c**). Consistent with these distinct modes of action, *uprA* overexpression restored LdtD-mediated β-lactam resistance in Δ*mltG* cells (**Fig. 4d**), whereas *mltG* overexpression only partially rescued the Δ*uprA* phenotype (**Fig. 4e**). Thus, UprA and MltG perform mechanistically distinct, yet partially overlapping functions in glycan-strand release.

### UprB acts on ECA and O-antigen lipidic precursors

Having established that UprA cleaves C55-linked peptidoglycan intermediates, we next asked whether other Upr-like proteins act on lipid-linked substrates from additional C55-dependent envelope pathways. We focused first on UprB, whose deletion was previously shown to cause a strong increase in ECA production^20^ (**Fig. 5a**), a phenotype that points to a direct connection with the ECA biosynthetic route. We therefore hypothesized that UprB might act on lipid-linked ECA precursors.

**Figure 5.**
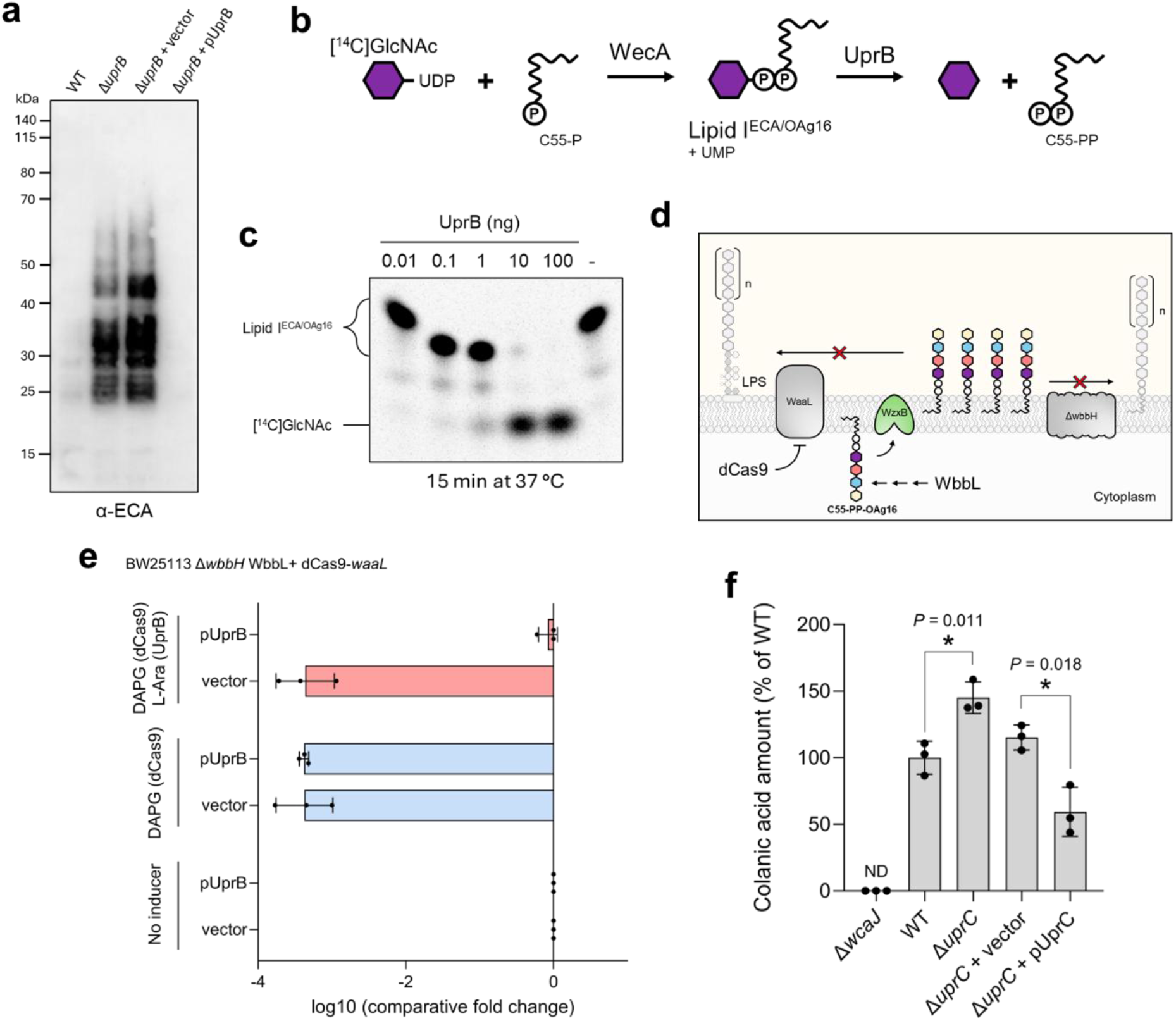
UprB acts on ECA and OAg16 precursors and UprC is linked to the colanic acid pathway. (**a**) Deletion of *uprB* leads to an increased production of enterobacterial common antigens (ECAs). ECAs were detected by Western blot. (**b**) Schematic illustrating the experimental procedure to test the *in vitro* activity of UprB on lipid I^ECA/OAg16^. [^14^C]-labelled UDP-GlcNAc was incubated with C55-P and WecA to generate lipid I^ECA/OAg16^. The product of the reaction was purified and incubated with UprB. (**c**) UprB hydrolyzes lipid I^ECA/OAg16^ *in vitro*. The products of the reaction were separated by thin layer chromatography and the radioactive signal of the [^14^C]GlcNAc was measured. (**d**) Schematic illustrating the experimental procedure to accumulate precursors of the OAg16 pathway *in vivo*. Production of OAg16 is restored by the expression of *wbbL*, polymerization of OAg16 is abolished by the deletion of *wbbH*, and the transfer of OAg16 on LPS is abolished by the repression of *waaL* by dCas9. (**e**) Plating efficiency of *E. coli* BW25113 Δ*wbbH* carrying plasmids for the expression of *wbbL*, dCas9 directed to *waaL*, and *uprB*. Growth efficiencies were determined relative to the growth of the strains in the absence of *waaL* repression by dCas9 (No inducer). BHI agar plates contained 40 µM IPTG, 50 µM DAPG, and 1% L-arabinose for induction of *wbbL*, *dCas9*, and *uprB*, respectively. The horizontal bars correspond to the mean, the error bars represent the standard deviation, and the dots represent individual data (n = 3 biological replicates). (**f**) Deletion of *uprC* leads to an increased production of colanic acids. Colanic acids were extracted from cultures grown at 22 °C in M9 medium for 3 days and quantified with the m-hydroxydiphenyl method^42,43^. The vertical bars correspond to the mean of the colanic acid amount relative to WT, the error bars represent the standard deviation, and the dots represent individual data (n = 3 biological replicates). ND, not detected.

To test this, we used a synthetic [^14^C]-labelled lipid I^ECA/OAg16^ as a simplified *in vitro* substrate. This minimal precursor consists of a single GlcNAc residue attached to C55 via a diphosphate linkage, the common initiating unit shared by both the ECA and O-antigen 16 (OAg16) biosynthetic pathways^9,10^ (**Fig. 5b**; **Extended Data Fig. 1b,c**). Strikingly, UprB efficiently cleaved this substrate, releasing the [^14^C]-labelled GlcNAc from the C55-PP lipid carrier (**Fig. 5c**). Because lipid I^ECA/OAg16^ is common to both the ECA and OAg16 pathways, this result raised the possibility that UprB also acts on OAg16 precursors *in vivo*. We therefore engineered a strain that accumulates OAg16 lipidic precursors in the periplasm, thereby sequestering the C55 lipid carrier. This strain expressed *wbbL* (**Fig. 5d**), encoding the rhamnosyltransferase required to restore OAg16 precursor synthesis^38^, in a Δ*wbbH* background lacking the periplasmic OAg16 polymerase^39^ (**Fig. 5d**; **Extended Data Fig. 1c**), thereby blocking polymer elongation in the periplasm (**Fig. 5d**). In parallel, *waaL*, encoding the ligase that transfers OAg16 to LPS and thereby releases C55-PP^40^ (**Fig. 5d**; **Extended Data Fig. 1c**), was downregulated by CRISPRi^41^ to trap OAg16 precursors on the lipid carrier (**Fig. 5d**). As expected, accumulation of OAg16 lipidic precursors resulted in a marked decrease in viability (**Fig. 5e**). Remarkably, overexpression of *uprB* fully restored growth (**Fig. 5e**). Together, these results indicate that UprB acts not only on ECA lipidic precursors but also on OAg16 lipidic intermediates, thereby releasing and redistributing the C55 lipid carrier to other envelope biogenesis pathways. Whether UprB also acts on polymerized, WaaL-competent OAg16 chains — for instance as a quality-control mechanism when ligation is delayed — remains an open question. Nevertheless, modeling of UprB with one- or three-subunit long ECA or OAg16 precursors shows that polymerized ECA or OAg16 chains can readily fit within the UprB active-site cavity and place the scissile phosphoacetal bond near the Thr^165^-His^189^-Arg^192^-Glu^240^ cluster (**Extended Data Fig. 6a-d**).

### UprC is linked to the colanic acid pathway

In addition to UprA and UprB, *E. coli* encodes a third periplasmic Upr-like protein, UprC (formerly YgjQ; **Extended Data Fig. 2**), whose function was also unknown. Given the activities defined above, we asked whether UprC, too, acts as a release factor for a C55-PP-linked polysaccharide intermediate. Loss of *uprC* causes neither peptidoglycan accumulation nor an increase in ECA production (**Extended Data Fig. 7a,b**), distinguishing UprC from both UprA and UprB. We therefore turned to the remaining C55-dependent pathway in *E. coli* — colanic acid synthesis (**Extended Data Fig. 1d**). Strikingly, direct quantification^42,43^ revealed that Δ*uprC* cells produce significantly more colanic acid than wild type (**Fig. 5f**). In addition, modeling positions C55-PP-colanic acid precursors within the catalytic cavity of UprC, with the scissile phosphoacetal bond oriented toward the conserved Thr^132^-His^156^-Arg^159^-Glu^191^ cluster (**Extended Data Fig. 8a,b**). Together, these observations identify UprC as the likely release factor for the colanic acid pathway, although direct biochemical validation on a defined colanic acid substrate is still needed.

### Upr-like proteins are widespread enzymes that share a common catalytic activity but partition clearly *in vivo*

Because UprA, UprB, and UprC share a conserved Upr-like domain, we asked whether they also exhibit overlapping substrate specificity. *In vitro*, all three enzymes hydrolyze purified lipid II^PG^ and release the same GlcNAc-MurNAc-pentapeptide product, as monitored by LC-MS (**Fig. 6a**). Thus, the conserved Upr-like fold defines a common catalytic activity — hydrolysis of the C55-PP-sugar bond — that is retained across the family. *In vivo*, however, the situation is sharply different. Loss-of-function phenotypes are pathway-specific: only Δ*uprA* causes peptidoglycan accumulation (**Extended Data Fig. 7a**), only Δ*uprB* increases ECA production (**Extended Data Fig. 7b**), and only Δ*uprC* elevates colanic acid levels (**Extended Data Fig. 7c**). Gain-of-function assays lead to the same conclusion: neither *uprB* nor *uprC* complements Δ*uprA* in the LdtD-mediated β-lactam resistance assay (**Fig. 6b**), and neither *uprA* nor *uprC* rescues the loss of viability caused by OAg16 precursor accumulation in Δ*uprB* cells (**Fig. 6c**).

**Figure 6.**
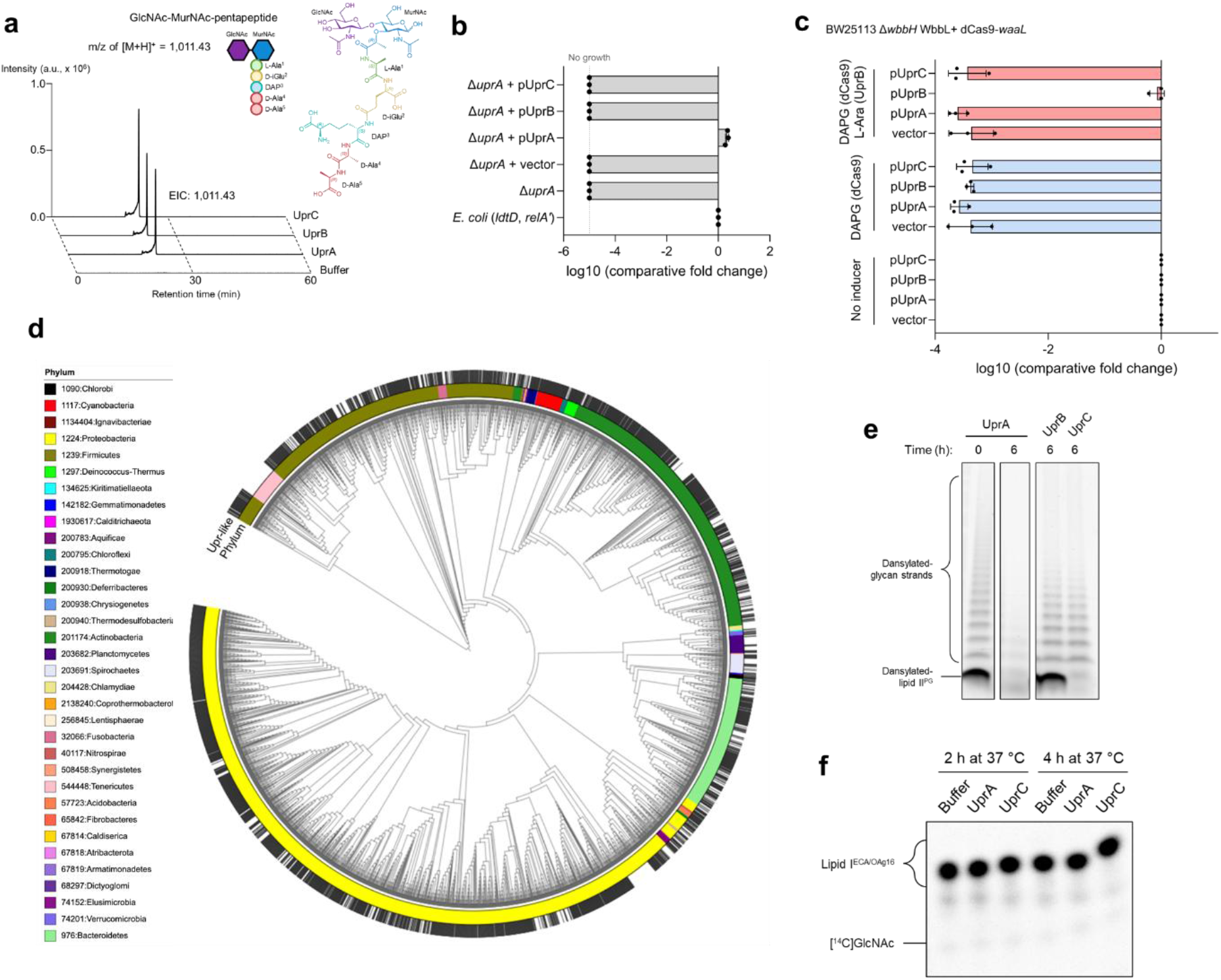
Upr-like proteins are widespread enzymes sharing a common catalytic activity but partition *in vivo*. (**a**) Extracted ion chromatograms (EICs) of the *in vitro* enzymatic reaction of UprA, UprB and UprC with purified lipid II^PG^. The structure of the detected product GlcNAc-MurNAc-pentapeptide is shown for reference. a.u., arbitrary unit. (**b**) Plating efficiency of *E. coli* (*ldtD*, *relA’*) and its derivatives harboring the deletion of *uprA* and plasmids for complementation with Upr-like enzymes. Growth efficiencies were determined relative to the growth of the parental strain *E. coli* (*ldtD*, *relA’*). Growth was tested in the presence of 2 µg/mL ceftriaxone, and 50 µM DAPG for induction of *uprA*, *uprB*, or *uprC*. BHI agar plates contained 40 µM IPTG and 1% L-arabinose for induction of *ldtD* and *relA’*, respectively. The horizontal bars correspond to the mean, the error bars represent the standard deviation, and the dots represent individual data (n = 3 biological replicates). (**c**) Plating efficiency of *E. coli* BW25113 Δ*wbbH* carrying plasmids for the expression of *wbbL*, dCas9 directed to *waaL*, and *uprA*, *uprB* or *uprC*. Growth efficiencies were determined relative to the growth of the strains in the absence of *waaL* repression by dCas9 (No inducer). BHI agar plates contained 40 µM IPTG, 50 µM DAPG, and 1% L-arabinose for induction of *wbbL*, *dCas9*, and *uprA*, *uprB* or *uprC*, respectively. The horizontal bars correspond to the mean, the error bars represent the standard deviation, and the dots represent individual data (n = 3 biological replicates). (**d**) Phylogenetic tree showing distribution of Upr-like proteins (black bars) in a diverse set of 3992 bacterial taxa. Upr-like proteins are found in all phyla except in the *Mycoplasma* (*Tenericutes*) and *Chlamydiae*. (**e**) UprB and UprC are not active on polymerized glycan strands. Fluorescence of synthesized dansylated-lipid II^PG^ polymerized into glycan strands by PBP1bγ S^510^A and incubated with UprA, UprB, or UprC. The dansyl moiety, attached to the third amino acid of the peptide stem, fluoresces in a hydrophobic environment such as DDM micelles. A decrease in fluorescence indicates release of the dansyl moiety from the micelles. Of note, although UprB is active on lipid II^PG^ (see panel a), it is not active on dansylated-lipid II^PG^. (**f**) UprA and UprC are not active on lipid I^ECA/OAg16^ *in vitro*. The products of the reaction were separated by thin layer chromatography and the radioactive signal of the [^14^C]GlcNAc was measured.

Phylogenetic analysis revealed that Upr-like proteins are widely distributed across bacterial phyla, present in nearly all major lineages and absent only in *Mycoplasma* (*Tenericutes*) and *Chlamydiae* (**Fig. 6d**). Within the family, Upr-like sequences fall into four distinct clades, all sharing the conserved Thr-His-Glu-Arg catalytic motif (**Extended Data Fig. 9a,b**): UprA and UprC group within a UprA/UprC-like clade, UprB defines the UprB-like clade, and two additional clades absent from *E. coli* complete the family — suggesting that Upr-like enzymes may serve C55-PP-dependent pathways beyond those characterized here.

What underlies this strong *in vivo* specificity, given the shared activity on lipid II^PG^? A closer look reveals that the apparent biochemical redundancy is more limited than it first appears. Although UprB and UprC hydrolyze purified lipid II^PG^, UprB did not hydrolyze dansylated-lipid II^PG^, and only UprA releases polymerized glycan strands from the C55 lipid carrier (**Fig. 6e**), identifying it as the only paralogue active on the extended peptidoglycan substrate tested here. Conversely, neither UprA nor UprC cleaves lipid I^ECA/OAg16^ (**Fig. 6f**), identifying UprB as the only paralogue active on the initiating ECA/O-antigen unit.

Together, these results show that Upr-like enzymes share a common catalytic core but have evolved distinct substrate preferences. These intrinsic preferences, likely combined with extrinsic factors such as subcellular localization, protein-protein interactions, and differential access to pathway-specific substrates, may account for the pathway partitioning observed *in vivo*, *i.e*. UprA, UprB, and UprC specifically act on peptidoglycan, ECA/O-antigen, and colanic acid, respectively.

## DISCUSSION

Here we identify the Upr-like proteins (formerly DUF218) as the first family of C55-PP-polysaccharide release factors. In *E. coli*, three periplasmic members — UprA, UprB, and UprC — hydrolyze the phosphoacetal bond that tethers envelope polysaccharide precursors to the C55 lipid carrier, liberating nascent polymers from the membrane and generating C55-PP for recycling. UprA acts on lipid II^PG^ (**Fig. 2 and 3**) and on elongating peptidoglycan glycan strands (**Fig. 4**), catalyzing the long-sought terminal step of peptidoglycan synthesis. UprB releases ECA and O-antigen precursors from C55 (**Fig. 5a-e**), while UprC limits the buildup of intermediates in the colanic acid pathway (**Fig. 5f**). Together, these activities define a unifying enzymatic principle: dedicated release factors that couple polysaccharide liberation to lipid-carrier recycling across bacterial envelope biogenesis pathways.

The identification of UprA resolves a longstanding puzzle in peptidoglycan biochemistry. Glycosyltransferases such as RodA, FtsW, and class A PBPs polymerize lipid II^PG^ into elongating glycan strands by transferring the growing polymer onto newly arrived lipid II^PG^ molecules, without forming a covalent enzyme-substrate intermediate^31,32^. The nascent strand therefore remains covalently tethered to C55-PP throughout elongation, and its release necessarily requires hydrolysis of the phosphoacetal bond — a reaction that had been postulated but whose enzymatic basis was unknown^44^. Our *in vitro* data demonstrate that UprA catalyzes precisely this reaction: it hydrolyzes the lipid-glycan linkage of lipid II^PG^ and of polymerized glycan strands, liberating the polymer from the membrane and regenerating C55-PP (**Fig. 2 and 3**). Isotope-labelling experiments in [^18^O]H_2_O revealed ^18^O incorporation on the GlcNAc-MurNAc-pentapeptide product (**Fig. 3e**), establishing that water attacks the sugar side of the phosphoacetal bond rather than on the phosphate side. This mechanism is fundamentally distinct from that of MltG, the only enzyme previously implicated in glycan-strand release. MltG is a lytic transglycosylase that cleaves the glycan backbone after the seventh disaccharide, liberating the chain with a terminal 1,6-anhydroMurNAc while leaving a committed elongation intermediate still attached to C55^36^. UprA, by contrast, severs the lipid-glycan bond itself, achieving arrest of the glycan strand elongation, polymer release, and carrier regeneration in a single step.

Beyond solving the missing release step, our data indicate that UprA also functions as a direct regulator of peptidoglycan output. The marked accumulation of peptidoglycan in Δ*uprA* cells shows that glycan-strand release is not merely a terminal processing event, but a key control point in the overall flux through the pathway. More broadly, the Upr-like family appears to regulate the balance between competing envelope biogenesis routes by controlling when lipid-linked intermediates are released and when the finite C55 pool is returned for reuse. In this view, Upr-like proteins are not only release factors in a mechanistic sense, but also regulators of pathway flux and carrier allocation across the bacterial envelope.

The same architecture and the same cluster of catalytic residues are found in UprB and UprC (**Extended Data Fig. 2; Extended Data Fig. 6**). The shared fold across the three paralogues implies that substrate selectivity is encoded in peripheral cavity features rather than in divergence at the catalytic core — a prediction directly testable by structure-guided mutagenesis. Notably, Upr-like proteins are widely distributed across bacteria, being detected in all phyla examined except *Mycoplasma* (*Tenericutes*) and *Chlamydiae* (**Fig. 6d**). *Mycoplasma* lack a cell wall altogether and have shed much of the corresponding envelope biogenesis machinery, whereas *Chlamydiae* do produce peptidoglycan^45^, suggesting that this lineage and other species lacking Upr-like enzymes may rely on an unrelated enzyme for C55-PP-polysaccharide release. This raises the possibility that distinct protein families can fulfill analogous functions. One candidate is the VanZ family: VanZ was originally identified in the vancomycin resistance operon of enterococci^46^, and a recent study showed that the yeast VanZ-family member Llp1 acts as a dolichol-linked oligosaccharide diphosphatase involved in quality control of N-glycosylation precursors^47^. Whether bacterial VanZ proteins similarly act as lipid II^PG^ diphosphatases remains to be demonstrated, but the functional parallel — a VanZ-family enzyme cleaving a polyprenol-diphosphate-linked glycan — is suggestive.

How does UprA regulate peptidoglycan synthesis *in vivo*? Uncontrolled hydrolysis of lipid II^PG^ during exponential growth would be counterproductive, given the high demand for this precursor^13^. Indeed, hydrolyzing free lipid II^PG^ as rapidly as it is produced would establish a futile cycle, dissipating the energetic and lipid-carrier investment of precursor synthesis without yielding peptidoglycan. We therefore propose that UprA acts primarily on elongating glycan strands rather than on free lipid II^PG^. In this model, UprA terminates the processive activity of peptidoglycan-synthesizing complexes by releasing the nascent chain from C55. Such termination could interrupt ongoing polymerization, promote complex disassembly, and redistribute enzymatic activity to sites where new synthesis is required. This model aligns with the observation that *uprA* deletion rescues the growth defects of several cell division mutants^21^: if UprA repeatedly terminates ongoing synthesis at the septum, the divisome would depend more heavily on these proteins to sustain peptidoglycan production — a dependence that is relieved when UprA is absent. Identifying the regulators that control UprA activity will be essential to understanding how it is integrated into the spatial and temporal coordination of peptidoglycan synthesis during the cell cycle.

Beyond peptidoglycan, our data demonstrate that the Upr-like family partitions across envelope biogenesis pathways. UprB contributes to ECA homeostasis through its role in lipid-carrier release, and previous work has shown that this regulation requires the chain-length regulator WzzE^20^. Interestingly, WzzE is also required for cyclic ECA production^9^, raising the possibility that UprB acts in concert with WzzE to direct circularization of ECA chains rather than simple hydrolysis — a question that warrants direct biochemical investigation. The activity of UprB on O-antigen lipidic precursors (**Fig. 5b-e**) likely serves a different purpose: because O-antigen is normally transferred to LPS by the WaaL ligase^40^, UprB’s cleavage of O-antigen intermediates would primarily redistribute C55-P to other pathways rather than regulate O-antigen assembly *per se*. A conceptually similar lipid carrier salvage mechanism was recently proposed in *Bacillus subtilis*, in which C55-P can be released from non-essential lipid-linked intermediates on the cytoplasmic side of the membrane to prioritize peptidoglycan synthesis^48^. UprC, in turn, presumably acts on colanic acid precursors (**Fig. 5f**). As the membrane-release step of colanic acid synthesis remains poorly characterized^11^, UprC represents a strong candidate for the release factor in this pathway, although formal *in vitro* validation is still needed.

The broad distribution of Upr-like proteins across the bacterial kingdom suggests that enzymatic release of lipid-linked glycans is a widespread step in envelope biogenesis, and that Upr-like enzymes provide an evolutionarily conserved catalytic mechanism for this step. Within the family, sequences resolve into four distinct clades — UprA/UprC-like, UprB-like, and two additional groups (clades 3 and 4) — all retaining the conserved Thr-His-Glu-Arg catalytic motif (**Fig. 6d; Extended Data Fig. 9a,b**). An important prediction of our model is that the presence of UprB- and UprC-like orthologues in a given species should correlate with whether that organism incorporates ECA, O-antigen, or colanic acid into its envelope. Systematic analysis of orthologue distribution across diverse bacterial lineages could test this prediction and potentially reveal Upr-like proteins dedicated to additional C55-dependent pathways not present in *E. coli*, such as wall teichoic acid synthesis in *Firmicutes* or glycosylation of envelope proteins.

An additional, non-mutually exclusive role for Upr-like proteins may be in quality control of envelope precursor trafficking. Lipid-linked intermediates that reach the periplasm but fail to polymerize — for instance, lipid I^PG^ or lipid I^ECA/OAg16^ that are flipped prematurely^49^ — would sequester C55 unproductively. Our observation that UprA hydrolyzes lipid I^PG^ (**Fig. 2e**) and that UprB cleaves lipid I^ECA/OAg16^ (**Fig. 5c**) indicates that Upr-like enzymes can clear such aberrant intermediates, recycling the carrier and preventing pathway congestion. This surveillance function would be particularly important under conditions of stress or enzyme fluctuation. It would also explain the apparent non-essentiality of Upr-like proteins under standard laboratory conditions: their contribution may become critical precisely when homeostasis is challenged, and may otherwise be buffered by parallel mechanisms such as the lytic transglycosylase MltG, which cleaves a different bond but achieves an overlapping outcome for the peptidoglycan pathway.

The convergence of multiple essential envelope pathways on a single, finite lipid carrier makes C55 recycling an attractive target for antibacterial intervention^50^. The precedent is clear: clovibactin, a recently discovered antibiotic, kills drug-resistant Gram-positive pathogens by targeting the diphosphate moiety of C55-PP-linked precursors^51^, and amphomycin similarly disrupts C55-P availability^52–54^. Inhibiting Upr-like enzymes would trap C55 in dead-end complexes with polysaccharide cargo, simultaneously blocking polymer release and starving competing pathways of lipid carrier. Because all three *E. coli* Upr-like proteins share a conserved catalytic domain, a single inhibitor targeting this domain could, in principle, disrupt multiple envelope pathways at once. Whether such inhibitors can be developed, and whether they would synergize with existing antibiotics that target lipid II^PG^ and peptidoglycan synthesis — such as vancomycin or β-lactams — remains to be explored. Finally, *E. coli* encodes a fourth Upr-like protein, YdcF, which localizes to the cytoplasm, lacks several residues conserved in the periplasmic family members, and binds S-adenosyl-L-methionine *in vitro*^55^; its substrate and function remain undefined, and we retain the YdcF designation pending further characterization.

In sum, our findings establish the Upr-like protein family as C55-PP-polysaccharide release factors and reveal a unifying enzymatic principle governing polysaccharide liberation and lipid-carrier recycling across bacterial envelope biogenesis. By identifying the enzyme that catalyzes the terminal step of peptidoglycan synthesis and by demonstrating analogous activities in the ECA/O-antigen and colanic acid pathways, this work fills a major gap in our understanding of how bacteria build and maintain their cell envelope. The challenge ahead is to elucidate the structural basis of substrate selectivity, the regulatory mechanisms that control Upr-like activity *in vivo*, and the therapeutic potential of targeting this conserved and widespread enzymatic family.

## METHODS

### Strains, plasmids, and growth conditions

All strains used in this study were derived from *E. coli* BW25113. Details regarding bacterial strains, plasmid origins, and their properties are provided in **Supplementary Table S1**. Bacteria were cultivated in brain heart infusion (BHI; Difco) or Miller’s lysogeny (LB; Difco) broth or agar plates and maintained at 37 °C unless otherwise specified. Liquid cultures were aerated by shaking at 200 rpm. To prevent plasmid loss, growth media were consistently supplemented with appropriate antibiotics: tetracycline (5 µg/mL) for plasmids derived from pHV6, chloramphenicol (20 µg/mL) for plasmids derived from pKT8, pOV86 or pFR56, and spectinomycin (100 µg/mL) for plasmids derived from pHV193 and pBAD42. For β-lactam susceptibility assays, ceftriaxone was used at concentrations of 2 µg/mL. Induction of the P*_trc_*, P*_araBAD_*, and P*_hlf_* promoters was performed with isopropyl β-D-1-thiogalactopyranoside (IPTG, 40 µM), L-arabinose (1%), and 2,4-diacetylphloroglucinol (DAPG, 50 µM), respectively. Gene deletions were introduced into *E. coli* BW25113 by P1 transduction^56^ of the Km^R^ cassette from selected Keio collection mutants^57^. Transductants were selected with kanamycin (50 µg/mL). Plasmids constructed in this study were generated using NEBuilder HiFi DNA Assembly (New England Biolabs), unless otherwise specified. Directed mutagenesis was used to introduce substitutions for the construction of plasmids expressing *uprA* variants.

### Plating efficiency assay

Bacteria were cultivated until they reached the late growth phase, indicated by an optical density at 600 nm (OD600) greater than 1.0. The OD600 of the cultures were then adjusted to 1.0, and a series of 10-fold serial dilutions ranging from 10^-1^ to 10^-6^ were prepared in BHI or LB broth. 5 µL of each diluted bacterial suspension were spotted onto BHI or LB agar plates. Plates were supplemented with various inducers and drugs according to the figure legends. The plates were incubated at 37 °C and imaged after 16 hours (without ceftriaxone) or 24 hours (with ceftriaxone).

### Immunoblotting

Bacteria were grown at 37 °C to the early exponential phase (OD600 from 0.05 to 0.1). When needed, expression of Upr-like proteins was induced by adding DAPG (50 µM), followed by incubation for 1 h at 37 °C to the late exponential phase (OD600 0.4 to 0.8). Cells were precipitated on ice by adding 900 µL of culture to 100 µL of 100% trichloroacetic acid, centrifuged, and washed with 1 mL ice-cold acetone. Pellets were resuspended in SDS sample loading buffer (60 mM Tris-HCl pH 6.8, 2% SDS, 10% glycerol, 0.1 mg/mL bromophenol blue), with protein amounts normalized by adding 100 µL of buffer for 1 unit of OD600. Samples were boiled for 10 min and loaded on NuPAGE Bis-Tris 4-12% gels. Immunoblotting was performed using anti-ECA primary antibodies (1:10,000), followed by a horseradish peroxidase-conjugated anti-rabbit secondary antibody (Sigma, 1:10,000). Detection was carried out with Immobilon Western Chemiluminescent HRP Substrates (Millipore) and imaged using an Amersham ImageQuant 800 Western blot imaging system (Cytiva).

### Fluorescence microscopy imaging

Bacteria were grown in LB at 37 °C to mid- to late-exponential phase (OD600 ∼0.4-0.8) in the presence of 100 µM HADA for ∼2 h to label newly synthesized peptidoglycan. Cells were harvested and washed three times in PBS to remove excess dye, then immobilized on a 1% agarose PBS pad. Fluorescence imaging was performed using a Nikon Ti2-E fully motorized inverted epifluorescence microscope equipped with a CFI Plan Apochromat λ DM 100x/1.45 oil objective, a Sola SEII FISH illuminator (Lumencor), and a Prime95B camera (Photometrics). HADA fluorescence was detected using appropriate filter sets (excitation ∼405 nm). Images were processed and analysed with ImageJ.

### Peptidoglycan quantification and analysis by LC-MS

Bacteria were grown in LB broth supplemented with the appropriate inducers and antibiotics at 37 °C to mid-exponential phase (OD600 ∼0.4-0.6). Cells were harvested by centrifugation and sacculi were extracted by boiling in 4% SDS for 45 min. Sacculi were collected by centrifugation at 20,000 x g for 20 min at 20 °C and washed five times with H_2_O. Purified sacculi were treated with 100 µg/mL trypsin in 20 mM Tris-HCl (pH 7.5) for 16 h at 37 °C, washed four times with H_2_O, incubated for 10 min at 98 °C to inactivate the remaining enzyme, and washed a final time with H_2_O.

For peptidoglycan quantification, sacculi were dried for 16 h at 60 °C in a beaker and weighed. For muropeptide LC-MS analysis, sacculi were digested with lysozyme (120 µM) in 50 mM ammonium bicarbonate (pH 7.4) with shaking at 1000 rpm for 16 h at 37 °C. For the *in vitro* UprA digestion assay on muropeptides, UprA (0.35 µM) was added during the lysozyme treatment. Insoluble debris was removed by centrifugation, and the soluble fraction containing the muropeptides was reduced with sodium borohydride at 5 mg/mL in 125 mM borate buffer (pH 9.0) for 45 min. The reaction was acidified to pH 3.0 with formic acid, lyophilized, and resuspended in mobile phase A (H_2_O, 0.1% formic acid). Muropeptides were separated by injecting 5 µL of sample on a C18 column (Hypersil GOLD aQ, 150 x 2.1 mm, 1.9 µm; Thermo Fisher Scientific) at 50 °C on an Agilent 1290 HPLC system. The flow rate was constant at 0.4 mL/min using mobile phase A and B (acetonitrile, 0.1% formic acid). An Agilent 6546 ion funnel mass spectrometer (MS) was used in the positive mode with an electrospray ionization (ESI) (voltage 3500 V, Nozzle voltage 1000 V, sheath gas 350 °C at 11 L/min, nebulizer pressure 35 psi and drying gas 300 °C at 8 L/min). Elution was performed with a linear gradient (0 to 20%) of mobile phase B applied between 10 and 80 min.

### Remazol Brilliant Blue assay

Purified peptidoglycan was labelled with Remazol Brilliant Blue as described previously^58,59^. Labelled peptidoglycan was incubated for 16 h at 37 °C with 35 μg/mL lysozyme or UprA. The undigested peptidoglycan was pelleted, and the released material was quantified by measuring dye absorbance at 600 nm.

### Selection of *uprA* suppressors mapping in *prc*

Suppressor mutants were selected from *E. coli* BW25113 Δ*relA* Δ*uprA* pKT2(*ldtD*) pKT8(*relA’*) strain. The selection procedure was independently carried out starting with independent colonies. Briefly, a fresh colony of *E. coli* BW25113 Δ*relA* Δ*uprA* pKT2(*ldtD*) pKT8(*relA’*) was inoculated in 3 mL BHI broth supplemented with tetracycline and chloramphenicol to counter-select loss of plasmids. Bacteria were grown overnight at 37 °C with shaking (200 rpm). Overnight cultures (100 µL) were plated on BHI agar supplemented with 8 µg/mL ceftriaxone, 40 µM IPTG, and 1% L-arabinose. Plates were incubated overnight at 37 °C. Individual suppressor colonies were isolated and subsequently grown in liquid culture. Genomic DNA was extracted from overnight cultures (Wizard DNA extraction kit, Promega) and sequenced by Illumina sequencing (Novogene). For Illumina sequencing, paired-end reads of 150 bp were generated using a NovaSeq6000 system (>50x coverage). Read processing and identification of the mutations were performed with the ‘breseq’ pipeline^60^ (v.0.35.5).

### Purification of UprA and UprC

The same procedure was applied for the purification of all UprA variants and UprC. *E. coli* C43(DE3) competent cells were transformed with the pET22b derivatives, encoding for a C-terminal twinstrep-tagged Upr protein. An overnight pre-culture was used to inoculate LB broth containing 200 µg/mL ampicillin, and the culture was grown at 37 °C to an OD600 of ∼0.8. Production of Upr protein was induced by adding 500 µM IPTG at 37 °C for 3 h. Cells were harvested by centrifugation, the pellet was resuspended in lysis buffer (20 mM Tris-HCl pH 8.0, 300 mM NaCl) supplemented with 50 µg/mL DNase I and EDTA-free protease inhibitor cocktail, and the cells were lysed by two passages through a French press at 15,000 psi. The lysate was clarified by centrifugation at 20,000 rpm for 40 min at 4 °C. The membrane pellet was resuspended in lysis buffer supplemented with 1% DDM and solubilized for 2 h at 4 °C on a tube-roller, followed by clarification by centrifugation. The supernatant was incubated with Strep-Tactin Sepharose resin (Cytiva) for 1 h at 4 °C, loaded onto a polypropylene column (Thermo Fisher Scientific), washed with purification buffer (20 mM Tris-HCl pH 8.0, 150 mM NaCl, 0.03% DDM), and eluted with purification buffer supplemented with 2.5 mM D-desthiobiotin. Elution fractions were pooled, concentrated using a 50 kDa molecular weight cut-off to ∼0.5-1.0 mL, and separated by size-exclusion chromatography on a Superdex 200 10/300 GL column (Cytiva) equilibrated in purification buffer at a flow rate of 1.0 mL/min. Fractions were collected and analysed by SDS-PAGE, and peak fractions containing Upr protein were pooled and concentrated using a 50 kDa molecular weight cut-off. Aliquots were flash-frozen in liquid nitrogen and kept at -80 °C. Protein concentration was determined by measuring UV absorbance with a NanoDropTM 2000 spectrophotometer (Thermo Fisher Scientific).

### Purification of UprB

*E. coli* C43(DE3) competent cells were transformed with the pETHis60-*uprB* plasmid, encoding for a C-terminal 6xHis-tagged UprB. An overnight pre-culture was used to inoculate LB broth containing 100 µg/mL ampicillin, and the culture was grown at 37 °C to an OD600 of ∼1. Production of UprB was induced by adding 1 mM IPTG at 22 °C for 16 h. Cells were harvested by centrifugation, the pellet was resuspended in lysis buffer (20 mM Tris-HCl pH 7.4, 200 mM NaCl, 10% glycerol) supplemented with 50 µg/mL DNase I and EDTA-free protease inhibitor cocktail, and the cells were lysed by three passages through a French press at 15,000 psi. The lysate was clarified by centrifugation at 20,000 rpm for 40 min at 4 °C. The membrane pellet was resuspended in lysis buffer supplemented with 2% DDM and solubilized for 2 h at 4 °C on a tube-roller, followed by clarification by centrifugation. The supernatant was incubated with Ni-NTA-agarose beads (Qiagen) for 1 h at 4 °C, loaded onto a polypropylene column (Thermo Fisher Scientific), washed with purification buffer (20 mM Tris-HCl pH 8.0, 150 mM NaCl, 0.02% DDM), and eluted with purification buffer supplemented with 200 mM imidazole. Elution fractions were pooled and dialyzed for 5 h at 4 °C against purification buffer. Aliquots were flash-frozen in liquid nitrogen and kept at -80 °C. Protein concentration was determined by measuring UV absorbance with a NanoDropTM 2000 spectrophotometer (Thermo Fisher Scientific).

### Purification of PBP1bγ S^510^A

*E. coli* C43(DE3) competent cells were transformed with the pET22b-twinstrep-*mrcBγ* S^510^A plasmid, encoding for a C-terminal twinstrep-tagged PBP1bγ S^510^A. An overnight pre-culture was used to inoculate LB broth containing 200 µg/mL ampicillin, and the culture was grown at 37 °C to an OD600 of ∼0.8. Production of PBP1bγ S^510^A was induced by adding 500 µM IPTG at 16 °C for 16 h. Cells were harvested by centrifugation, the pellet was resuspended in lysis buffer (20 mM Tris-HCl pH 8.0, 300 mM NaCl) supplemented with 50 µg/mL DNase I and EDTA-free protease inhibitor cocktail, and the cells were lysed by two passages through a French press at 15,000 psi. The lysate was clarified by centrifugation at 20,000 rpm for 40 min at 4 °C. The membrane pellet was resuspended in lysis buffer supplemented with 1% DDM and solubilized for 2 h at 4 °C on a tube-roller, followed by clarification by centrifugation. The supernatant was incubated with Strep-Tactin Sepharose resin (Cytiva) for 1 h at 4 °C, loaded onto a polypropylene column (Thermo Fisher Scientific), washed with purification buffer (20 mM Tris-HCl pH 8.0, 150 mM NaCl, 0.03% DDM), and eluted with purification buffer supplemented with 2.5 mM D-desthiobiotin. Elution fractions were pooled, concentrated using a 50 kDa molecular weight cut-off to ∼0.5-1.0 mL, and separated by size-exclusion chromatography on a Superdex 200 10/300 GL column (Cytiva) equilibrated in purification buffer at a flow rate of 1.0 mL/min. Fractions were collected and analysed by SDS-PAGE, and peak fractions containing PBP1bγ S^510^A were pooled and concentrated using a 50 kDa molecular weight cut-off. Aliquots were flash-frozen in liquid nitrogen and kept at -80 °C. Protein concentration was determined by measuring UV absorbance with a NanoDropTM 2000 spectrophotometer (Thermo Fisher Scientific).

### Lipid II^PG^ extraction and purification

Lipid II^PG^ extraction was performed as previously described^61^. Briefly, the strain of *E. coli* BW25113 Δ*murJ* pHV193-*murJ* A^29^C expressing the MurJ A^29^C variant was grown in 1.5 L of LB at 37 °C until an OD600 of 0.4-0.6. Accumulation of lipid II^PG^ was triggered by adding 200 µM sodium (2-Sulfonatoethyl)methanethiosulfonate (also known as MTSES) for 10 min to inhibit MurJ A^29^C. Cells were centrifuged for 10 min, and the pellet was resuspended in 50 mM Tris pH 7.5. The lipids were extracted by mixing the cell suspension with 52.5 mL of CHCl_3_:MeOH (1:2, v:v) for 1 h at room temperature with strong agitation. The mixture was centrifuged and the supernatant was mixed to 30 mL of CHCl_3_ and 22.5 mL of 50 mM Tris pH 7.5 for 1 h at room temperature with strong agitation. The mixture was centrifuged and the interface containing the lipid II^PG^ was collected and lyophilized. The lyophilized interface was resuspended in 7.5 mL of 6 M pyridinium acetate:n-butanol (1:2, v:v) and 7.5 mL of n-butanol saturated H_2_O. The mixture was vortexed for 1 min, centrifuged, and the organic phase (upper phase) was collected. The aqueous phase (lower phase) was mixed with 5 mL of 6 M pyridinium acetate:n-butanol (1:2, v:v), vortexed, centrifuged, and the organic phase was collected and combined to the previous organic phase. The pooled organic phases were washed three times by the addition of 5 mL of n-butanol-saturated H_2_O, vortexing, centrifugation, and removal of the aqueous phase. The washed n-butanol phase containing the lipid II^PG^ was dried under a nitrogen flux, and the lipid II^PG^ was resuspended in 150 µL DMSO. To check the presence of lipid II^PG^, 5 µL of purified lipid II^PG^ in DMSO were mixed with 40 µL H_2_O and 5 µL of 100 mM ammonium acetate pH 4.2. The lipid tail of lipid II^PG^ was removed by heating at 98 °C for 30 min, and the product was lyophilized. The lyophilized product was resuspended in H_2_O supplemented with formic acid 0.1% and analysed by LC-MS to identify the phosphate-GlcNAc-MurNAc-pentapeptide resulting from the acidic hydrolysis of lipid II^PG^. LC-MS analysis was performed using the same procedure as described above for muropeptide analysis.

### Lipid substrate synthesis

His-tagged proteins MraY from *Aquifex aeolicus*, MurG from *E. coli* and WecA from *Thermotoga maritima* were prepared as previously described and were used for lipid substrates synthesis^62–65^.

The C55-PP-MurNAc-pentapeptide-GlcNAc (lipid II^PG^) synthesis was carried out in a volume of 200 µL containing 100 mM Tris-HCl pH 7.4, 150 mM NaCl, 40 mM MgCl_2_, 0.75% Triton X100, 1 mM C55-P, 1 mM UDP-MurNAc-pentapeptide, and 1.1 mM UDP-GlcNAc. The dansylated-lipid II^PG^ synthesis was carried out by replacing the UDP-MurNAc-pentapeptide with a dansyl-labeled UDP-MurNAc-pentapeptide. The reactions were initiated by adding MraY and MurG enzymes (20 µg each) and incubated for 4 h at 37 °C with shaking. The lipid II^PG^ products were purified by butanol extraction. Briefly, one volume reaction of 6 M pyridinium acetate and two volume reaction of 1-butanol were added to the mixtures, mixed by vortex and incubated for 2 h at -20 °C. The butanol phase was recovered by centrifugation for 5 min at 10 000 g and the lipid II^PG^ and dansylated-lipid II^PG^ concentrations were determined by amino acid analysis of a sample after hydrolysis in 6 M HCl for 16 h at 95 °C.

The C55-PP-GlcNAc (lipid I^ECA/OAg16^) synthesis was carried out in a volume of 200 µL containing 100 mM Tris-HCl pH 8.0, 10 mM MgCl_2_, 5% Triton X100, 1 mM C55-P, 1.1 mM UDP-GlcNAc and 120 µM radiolabeled UDP-[^14^C]GlcNAc. The reaction was initiated with the addition of WecA enzyme (20 µg) and the mixture was incubated for 8 h at 60 °C. The C55-PP-GlcNAc product was purified by butanol extraction as reported above. The C55-PP-GlcNAc concentration was determined by radioactivity measurement.

### Hydrolysis *in vitro* assay on lipid II^PG^

Hydrolysis assays were performed in a 10 µL reaction mixture containing 50 mM HEPES pH 7.5, 150 mM NaCl, 10 mM MgCl_2_, 10 mM CaCl_2_, 0.05% DDM, and 50 µM lipid II^PG^ (purified or synthesized) or dansylated-lipid II^PG^ (synthesized). Assays with purified lipid II^PG^ contained 10% DMSO. The reaction was initiated with the addition of Upr-like enzyme, and the mixture was incubated for 16 h at 37 °C. The reaction was stopped by heating at 95 °C for 2 min. For hydrolysis assay on lipid II^PG^, the substrates and products were separated by injecting 5 µL of sample on a C18 column (Hypersil GOLD aQ, 150 x 2.1 mm, 1.9 µm; Thermo Fisher Scientific) at 50 °C on an Agilent 1290 HPLC system. The flow rate was constant at 0.4 mL/min using mobile phase A and B (acetonitrile, 0.1% formic acid). An Agilent 6546 ion funnel mass spectrometer (MS) was used in the positive mode with an electrospray ionization (ESI) (voltage 3500 V, Nozzle voltage 1000 V, sheath gas 350 °C at 11 L/min, nebulizer pressure 35 psi and drying gas 300 °C at 8 L/min). Elution was performed with a linear gradient (0 to 20%) of mobile phase B applied between 10 and 80 min. For hydrolysis assay on dansylated-lipid II^PG^, 2.5 µL of SDS sample loading buffer 5X was added to the mixture, and the substrates and products were separated by SDS-PAGE in a 16% tricine precast gel (Thermo Fisher Scientific) at 125 V for 2 h. Fluorescence was imaged on an Amersham ImageQuant 800 Western blot imaging system (Cytiva) running on the UV (365 nm) channel.

### *In vitro* hydrolysis assay on dansylated-glycan strands

Dansylated-glycan strands were synthesized in a 9 µL reaction mixture containing 40 mM Tris-HCl pH 7.4, 150 mM NaCl, 10 mM MgCl_2_, 10 mM CaCl_2_, 0.1% DDM, and 50 µM dansylated-lipid II^PG^. The reaction was initiated with the addition of 0.35 µM PBP1bγ S^510^A, and the mixture was incubated for 16 h at 25 °C. The reaction was stopped by heating at 95 °C for 2 min. After cooling at room temperature, Upr-like enzymes were added to the dansylated-glycan strands and further incubated for 0, 3, or 6 h at 37 °C. The reaction was stopped by heating at 95 °C for 2 min. 2.5 µL of SDS sample loading buffer 5X was added to the mixture, and the substrates and products were separated by SDS-PAGE in a 16% tricine precast gel (Thermo Fisher Scientific) at 125 V for 2 h. Fluorescence was imaged on an Amersham ImageQuant 800 Western blot imaging system (Cytiva) running on the UV (365 nm) channel.

### *In vitro* hydrolysis assay on radiolabeled lipid I^ECA/OAg16^

Hydrolysis assays were performed in a 10 µL reaction mixture containing 40 mM Tris-HCl pH 7.4, 150 mM NaCl, 10 mM MgCl_2_, 10 mM CaCl_2_, 0.1% DDM, and 50 µM radiolabeled [^14^C]-lipid I^ECA/OAg16^. The reaction was initiated with the addition of Upr-like enzyme, and the mixture was incubated for 15 min at 37 °C (for UprB) or 2 h at 37 °C (for UprA and UprC). The reaction was stopped by freezing in liquid nitrogen. The substrates and products were separated by TLC on Silica gel 60 coated plates (Merck) using 1-propanol/ammonium hydroxide/H_2_O (6:3:1, v/v/v) as the mobile phase. Radioactive profile was determined and analysed using the Rita Star radioactivity detector for TLC (Elysia Raytest).

### Colanic acid extraction and quantification

Derivatives of *E. coli* BW25113 pOV86-*rcsA* were grown in M9 minimal medium supplemented with chloramphenicol (20 µg/mL) to counter-select loss of plasmid and 40 µM IPTG to induce the expression of *rcsA*. Cultures were incubated for 3 days at 22 °C with shaking at 200 rpm, except for experiments containing the Δ*uprB* strain (Extended Data Fig. 7c), which were grown at 30 °C as Δ*uprB* does not grow at 22 °C. For each sample, 5 mL of culture was lysed in a water bath at 100 °C for 15 min. After cooling to room temperature, the sample was centrifuged at 12,000 g for 10 min and the supernatant was collected. The supernatant was mixed with three volumes of ice-cold ethanol and incubated at 4 °C for 16 h. The precipitate was collected by centrifugation at 10,000 g for 15 min, dried, and resuspended in 1 mL H_2_O. Colanic acid content was quantified by measuring uronic acids using the m-hydroxydiphenyl method^42,43^. Briefly, the resuspended precipitate was mixed with 5 mL of H_2_SO_4_ supplemented with 25 mM sodium tetraborate and heated at 100 °C for 5 min. After cooling to room temperature, 200 µL of 1.5 mg/mL m-hydroxydiphenyl were added to the sample. Absorbance was measured at 526 nm, and colanic acid concentration was determined from a glucuronic acid standard curve.

### Bioinformatic analysis

#### AI co-folding calculations

Protein-ligand complexes of UprA, UprB, and UprC from *E. coli* with ligands were generated using Protenix web server (https://protenix-server.com) with the default parameters^26^. Protein sequences and ligand SMILES strings were provided as inputs. The generated structures were visualized with ChimeraX^66^.

#### Bacterial “Tree of life”

The reference genome was retrieved for each species available in the RefSeq database (accessed on April 14th, 2022) having a taxonomy defined in the NCBI’s Taxonomy Browser^67^, thus constituting a dataset of 3992 diverse and representative bacterial genomes. These were used as input for PhyloPhlAn version 3.2.1^68^ to generate the phylogenetic tree, using the databases PhyloPhlAn^69^ (with 400 universal marker genes), with the parameters --fast and --diversity high.

#### Phylogenetic tree of Upr-like protein sequences

Starting from the sequences of UprA, UprB, and UprC from *E. coli*, an HMM profile was generated using HMMER version 3.4 (http://hmmer.org/), then used to search for homologs of UprA, UprB, and UprC in the dataset of 3992 diverse genomes mentioned above. The resulting sequences were aligned using Clustal Omega version 1.2.4^70^ and manually filtered to conserve only sequences with conserved residues that are interacting directly with the ligands in the complexes generated by AI co-folding. The filtered sequences were realigned and the corresponding phylogenetic tree generated with Clustal Omega.

#### Phylogenetic trees figures

The phylogenetic trees figures were generated using iTOL^71^.

### Statistics and reproducibility

For the plating efficiency assays, and for peptidoglycan analysis and quantification assays, at least three biological repeats were performed. For *in vitro* hydrolysis assays with Upr-like enzymes, at least two technical repeats were performed. *P*-values were obtained from unpaired two-tailed *t*-test using Welch’s correction for heteroscedasticity.

## ACKNOWLEDGMENTS

We are indebted to members of the Collet lab for helpful suggestions and discussions. We thank Renato Morona (University of Adelaide) for the kind gift of anti-ECA antibody. This work was funded by Fonds de la Recherche Scientifique FRS-FNRS (grants WELBIO X.1509.24 and ARC 22/27-128 to J-FC). HV was supported by an EMBO postdoctoral fellowship (ALTF 359-2023) and an FRS-FNRS Chargé de Recherche postdoctoral fellowship. OV was supported by a FRIA-FNRS doctoral fellowship. TT, RA, and DP were supported by the French National Research Agency ANR “BacWall” (grant ANR-20-CE44-0009-01 to TT). BII was supported by the French National Research Agency ANR “Seq2Diag PPR Antibioresistance” (grant ANR-20-PAMR-0010 to BII).

## AUTHOR CONTRIBUTIONS

HV initiated the study; HV and J-FC supervised the study; HV, OV, RA, and DP performed the experiments; JJ provided technical assistance with the LC-MS setup; HV performed structural modeling; BII performed the phylogenetic analysis; HV, OV, RA, GTB, MA, J-EH, BII, TT, and J-FC analyzed the data; HV and J-FC wrote the paper.

## SUPPLEMENTARY INFORMATION

Supplementary Information is available for this paper.

## MATERIALS & CORRESPONDENCE

Correspondence and requests for materials should be addressed to

---

**Extended Data Figure 1.**
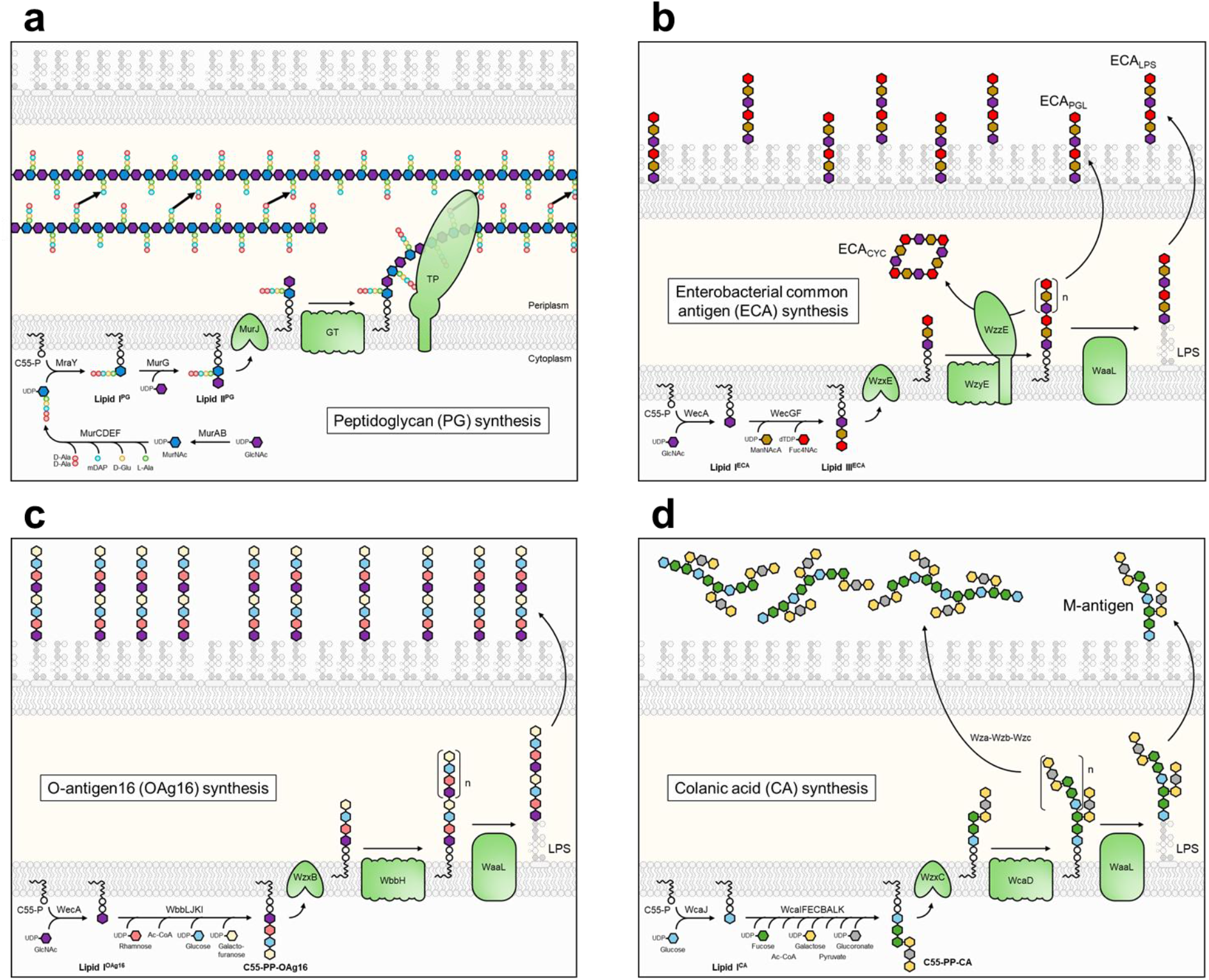
Envelope polysaccharide synthesis pathways relying on the C55-P lipid carrier in *Escherichia coli*. (**a**) Peptidoglycan (PG) synthesis pathway. The precursor is synthesized in the cytoplasm, linked to the C55-P lipid carrier, flipped to the periplasm, polymerized by the glycosyltransferases (GT) and cross-linked by the transpeptidases (TP). (**b**) Enterobacterial common antigen (ECA) synthesis pathway. The precursor is synthesized in the cytoplasm, linked to the C55-P lipid carrier, flipped to the periplasm, and polymerized by WzyE. ECA are circularized in the periplasm by an unknown enzyme dependent on the presence of WzzE (ECA_CYC_), linked to the LPS by the WaaL ligase (ECA_LPS_), or linked to a phosphatidylglycerol by an unknown enzyme and sent to the outer membrane (ECA_PGL_). (**c**) O-antigen 16 (OAg16) synthesis pathway. This pathway is cryptic in *E. coli* K-12 strains due to the insertion of an IS5 element in the *wbbL* gene. OAg16 synthesis can be restored by the introduction of an intact copy of *wbbL*. The precursor is synthesized in the cytoplasm, linked to the C55-P lipid carrier, flipped to the periplasm, polymerized by WbbH, and linked to LPS by the WaaL ligase. (**d**) Colanic acid (CA) synthesis pathway. The precursor is synthesized in the cytoplasm, linked to the C55-P lipid carrier, flipped to the periplasm, and polymerized by WcaD. CA are linked to the LPS by the WaaL ligase (M-antigen) or sent to the external medium by an elusive mechanism through the Wza-Wzb-Wzc complex. Abbreviations: GlcNAc, *N*-acetylglucosamine; MurNAc, *N*-acetylmuramic acid; ManNAcA, *N*-acetyl-D-mannosaminuronic acid; Fuc4NAc, 4-acetamido-D-fucose.

**Extended Data Figure 2.**
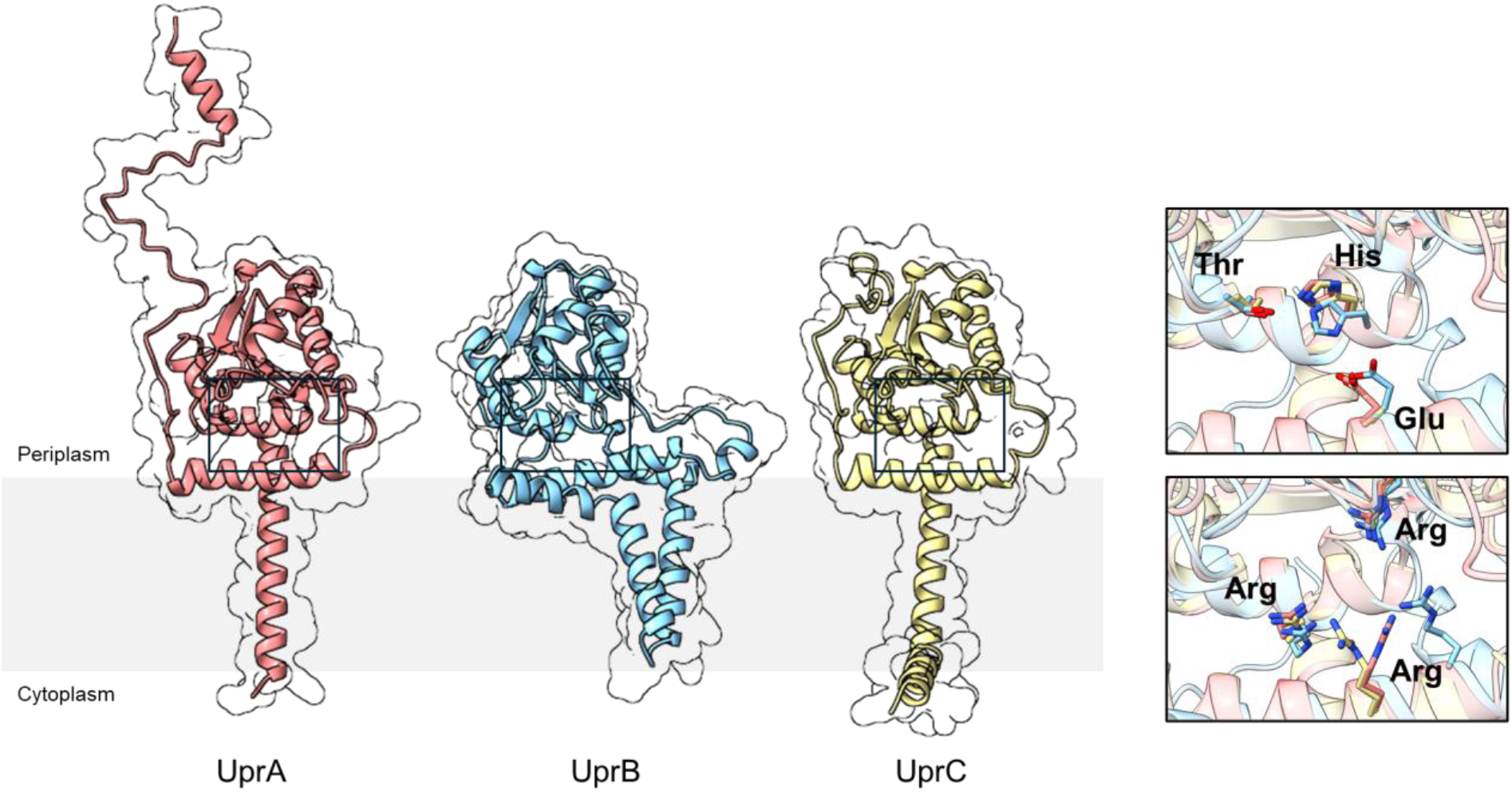
Upr-like proteins share conserved residues within a cavity. Structural comparison of predicted structures of UprA, UprB, and UprC from *E. coli*. The boxes show the conserved Thr, His, Glu, and Arg residues.

**Extended Data Figure 3.**
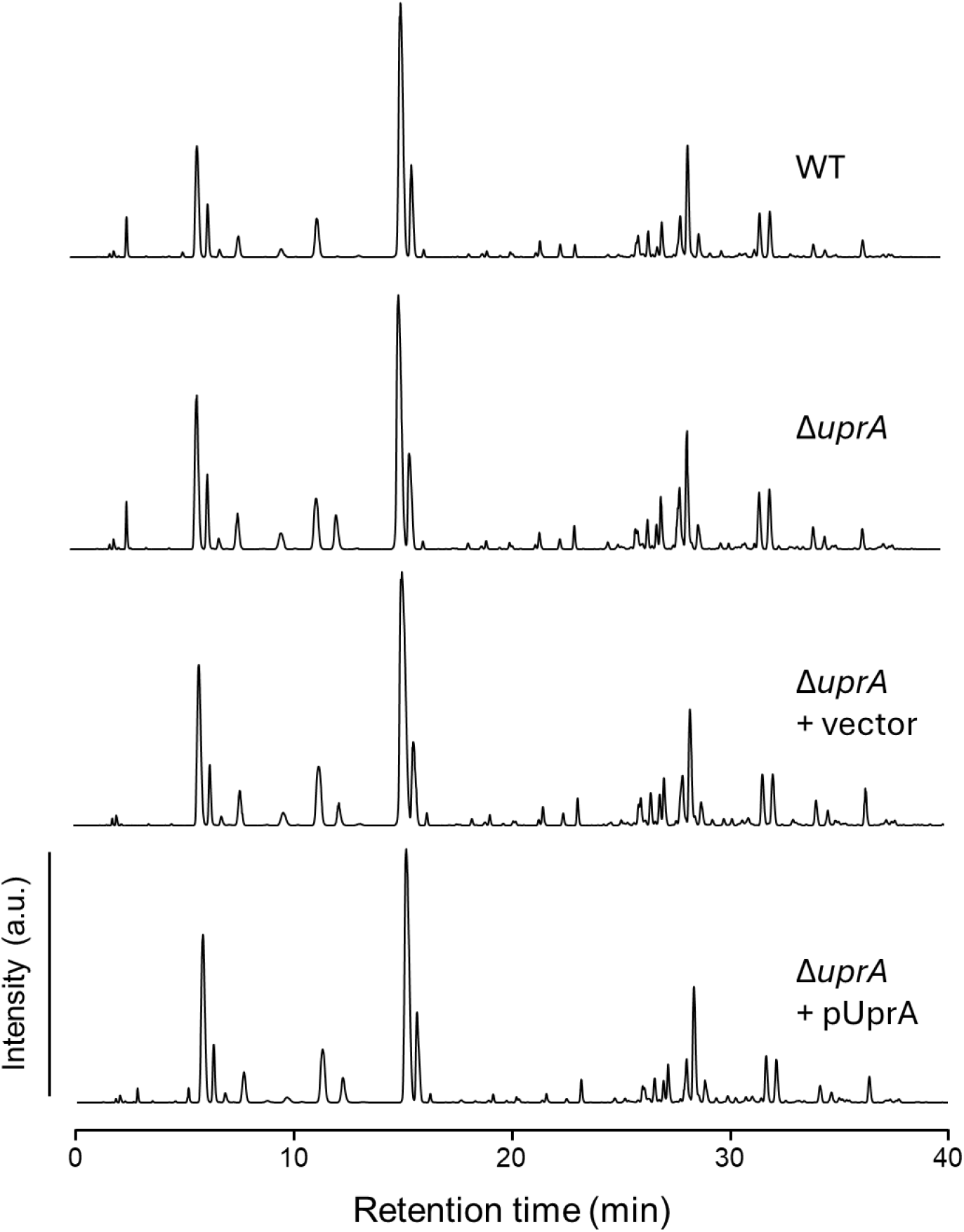
UprA deletion has no effect on the composition of peptidoglycan. Representative LC-MS profiles of muropeptides extracted from *E. coli* BW25113 (WT) and its derivatives harboring the deletion of *uprA* and plasmids for complementation. a.u., arbitrary unit.

**Extended Data Figure 4.**
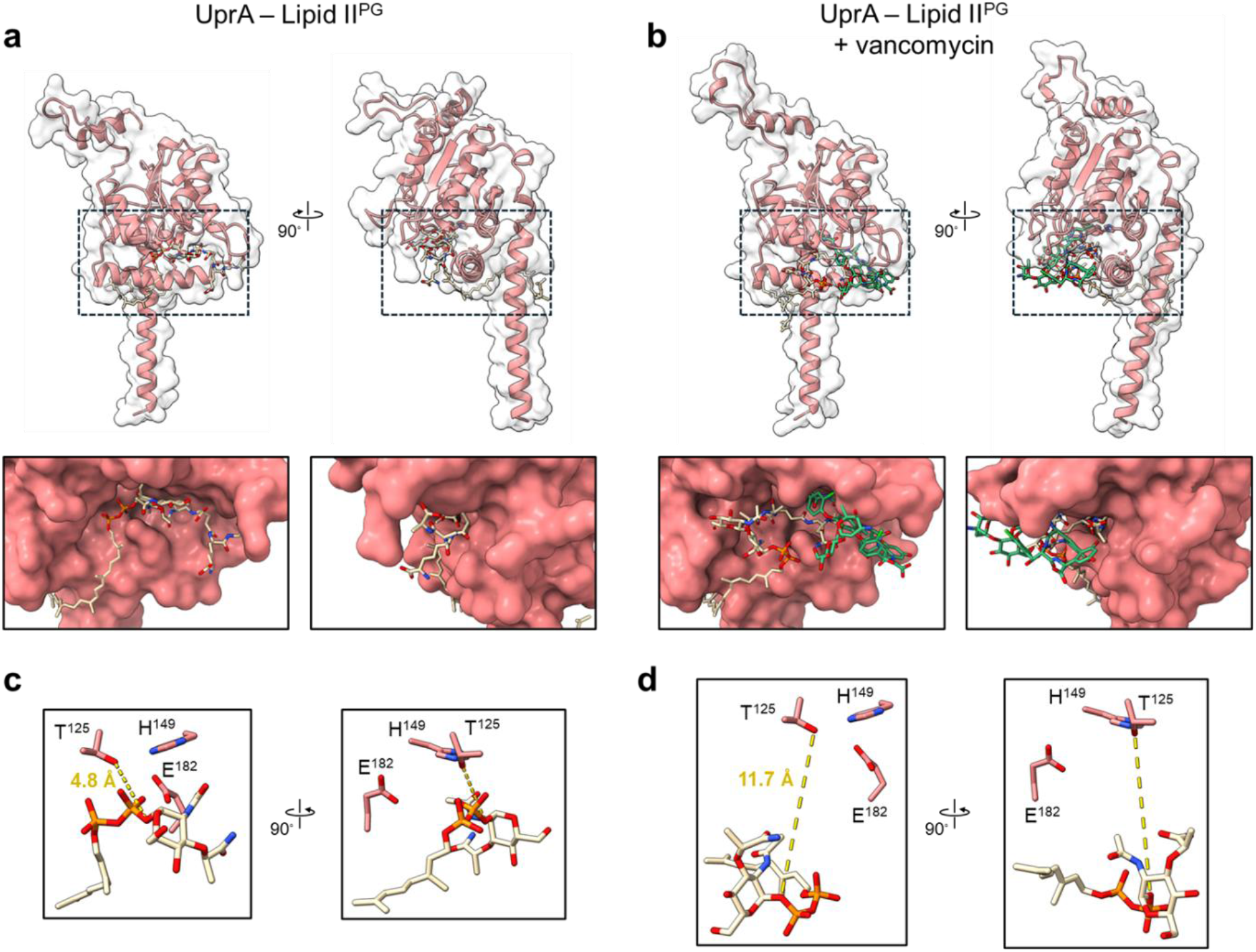
Modeling of UprA with lipid II^PG^ and vancomycin. (**a, b**) Modeling of UprA in complex with lipid II^PG^ without (**a**) and with (**b**) vancomycin. The boxed area shows a close-up view of the catalytic cavity of UprA. (**c, d**) Stick representation of the conserved residues T^125^, H^149^ and E^182^ with the lipid II^PG^ in the absence (**c**) or presence (**d**) of vancomycin. For the sake of simplicity, only the C10-PP-MurNAc part of lipid II^PG^ is shown. The distance between the hydroxyl function of T^125^ and the scissile phosphoacetal bond of the lipid II^PG^ is modelled at 4.8 Å in the absence of vancomycin and at 11.7 Å in the presence of vancomycin.

**Extended Data Figure 5.**
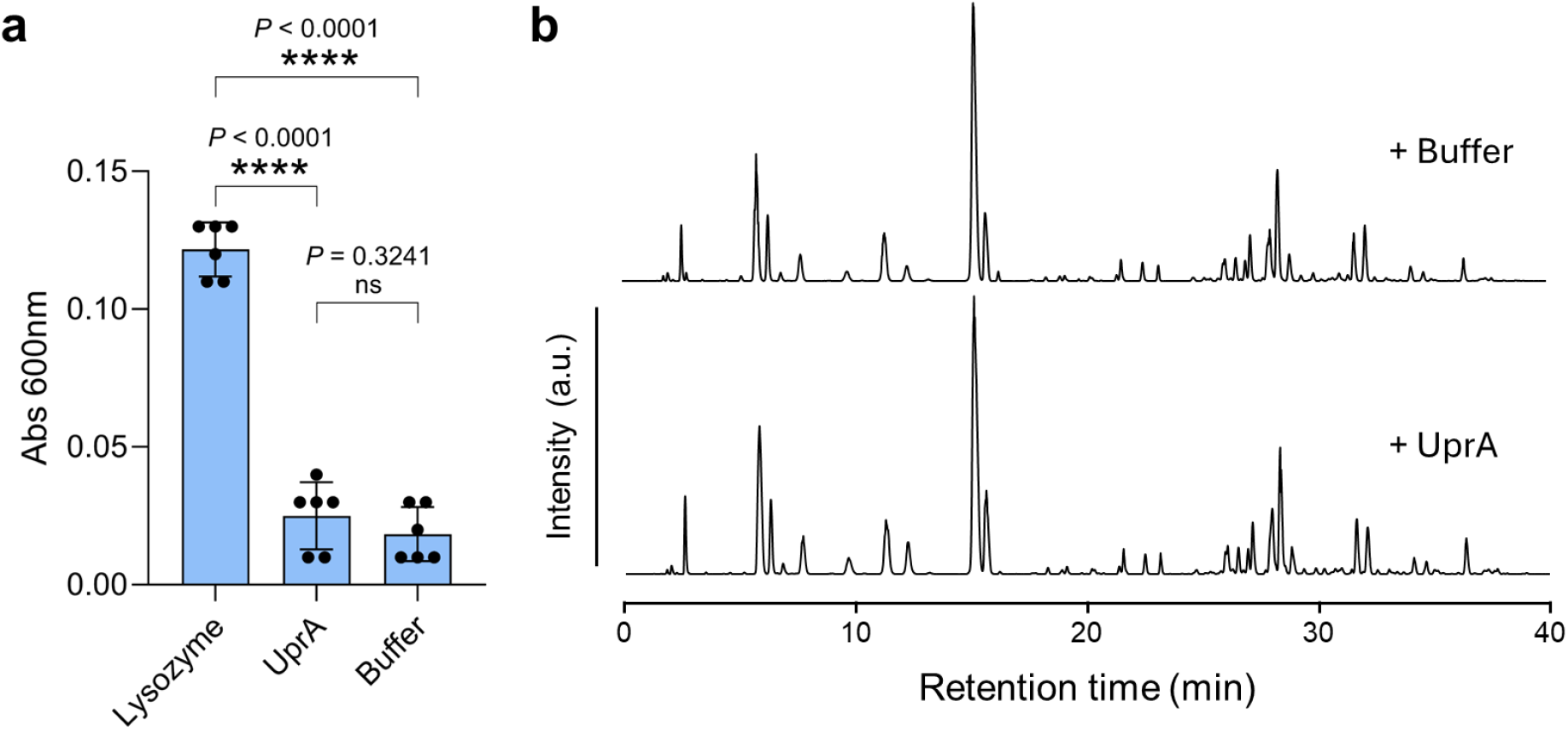
UprA is not a lytic transglycosylase, a muramidase, a glucosaminidase, an endopeptidase, or a carboxypeptidase. (**a**) Remazol brilliant blue-labeled peptidoglycan was incubated at 37 °C for 16 h with 35 μg/mL lysozyme or UprA. The undigested peptidoglycan was pelleted, and released material was quantified by measuring dye absorbance at 600 nm. The horizontal bars correspond to the mean of the absorbance, the error bars represent the standard deviation, and the dots represent individual data (n = 6 technical replicates). (**b**) Representative LC-MS profiles of muropeptides extracted from *E. coli* BW25113 (WT) and incubated with UprA. a.u., arbitrary unit.

**Extended Data Figure 6.**
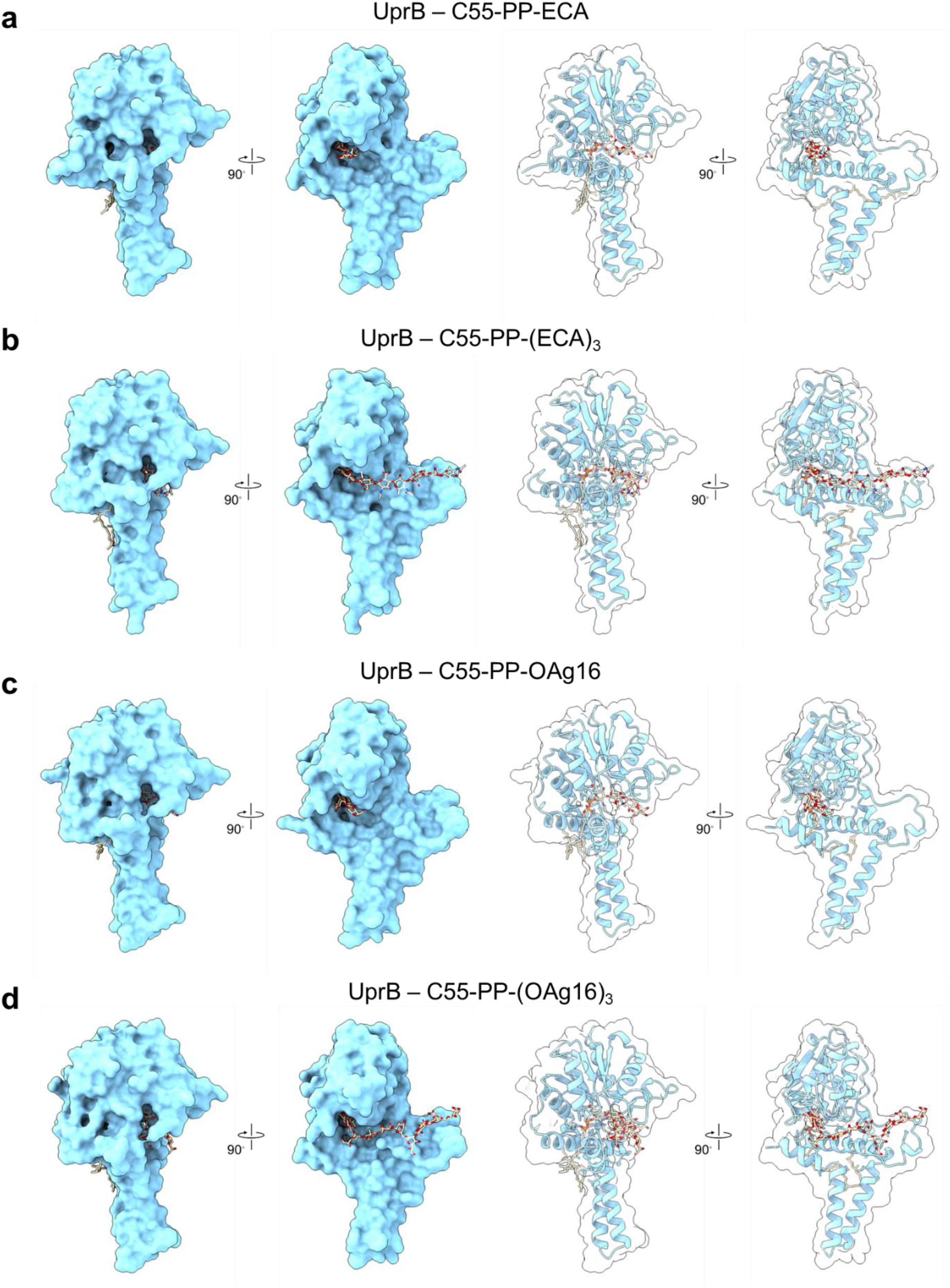
Modeling of UprB with lipidic precursors of the enterobacterial common antigen (ECA) and O-antigen 16 (OAg16) pathways. (**a, b, c, d**) Modeling of UprB with lipid III^ECA^ (**a**), with a three-subunit long lipid-linked ECA (**b**), with C55-PP-OAg16 (**c**), and with a three-subunit long lipid-linked OAg16 (**d**). All predictions show the substrate inside the catalytic cavity of UprB, with the scissile phosphoacetal bond oriented near the conserved catalytic residues.

**Extended Data Figure 7.**
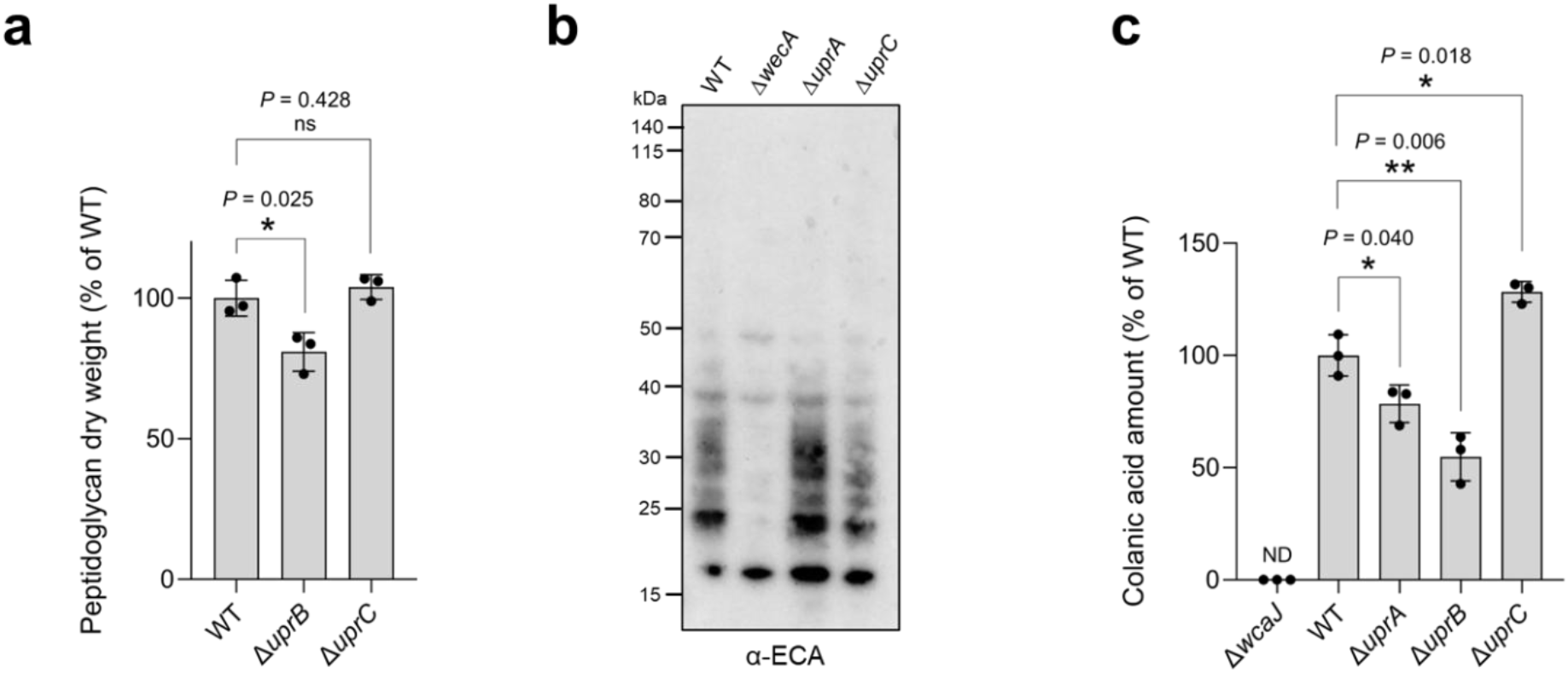
Upr-like enzymes are not redundant *in vivo*. (**a**) Dry weight quantification of peptidoglycan pellets of *E. coli* BW25113 (WT) and derivatives. In the tested conditions, *uprB* deletion leads to a decrease in peptidoglycan amount and the deletion of *uprC* has no effect. The vertical bars correspond to the mean of the peptidoglycan dry weight relative to WT, the error bars represent the standard deviation, and the dots represent individual data (n = 3 biological replicates). (**b**) The deletion of *uprA* or *uprC* has mild to no effect on the production of enterobacterial common antigens (ECAs). Δ*wecA* strain is the negative control that does not produce ECA. ECAs were detected by Western blot. (**c**) Colanic acids were extracted from cultures grown at 30 °C for 3 days in M9 medium and quantified with the m-hydroxydiphenyl method^42,43^. The deletion of *uprC* leads to an increase in colanic acids amounts, and the deletions of *uprA* or *uprB* both lead to a decrease in colanic acid amounts. The vertical bars correspond to the mean of the colanic acid amount relative to WT, the error bars represent the standard deviation, and the dots represent individual data (n = 3 biological replicates).

**Extended Data Figure 8.**
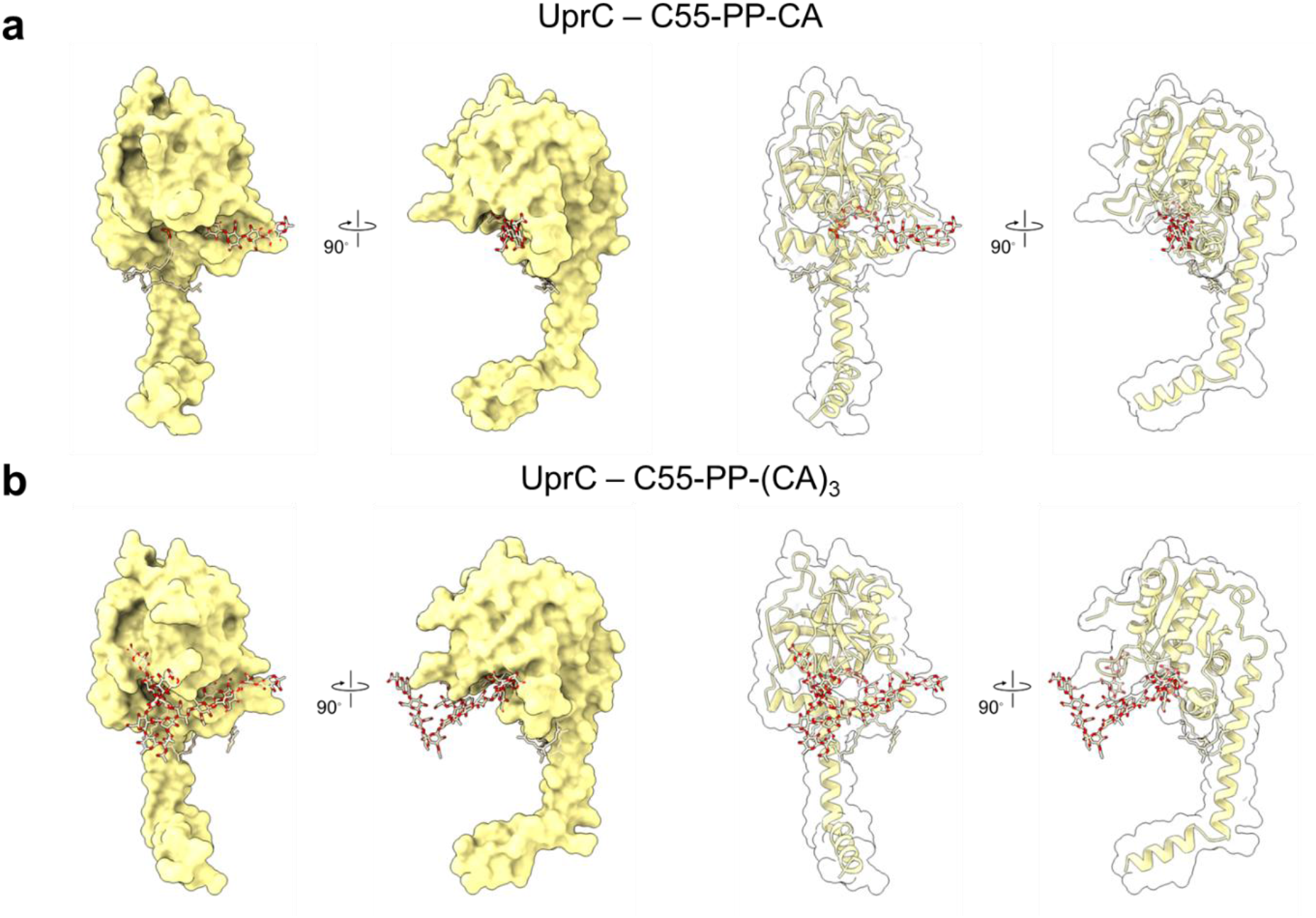
Modeling of UprC with lipidic precursors of the colanic acid (CA) pathway. (**a, b**) Modeling of UprC with C55-PP-CA (**a**), and with a three-subunit long lipid-linked CA (**b**). All predictions show the substrate inside the catalytic cavity of UprC, with the scissile phosphoacetal bond oriented near the conserved catalytic residues.

**Extended Data Figure 9.**
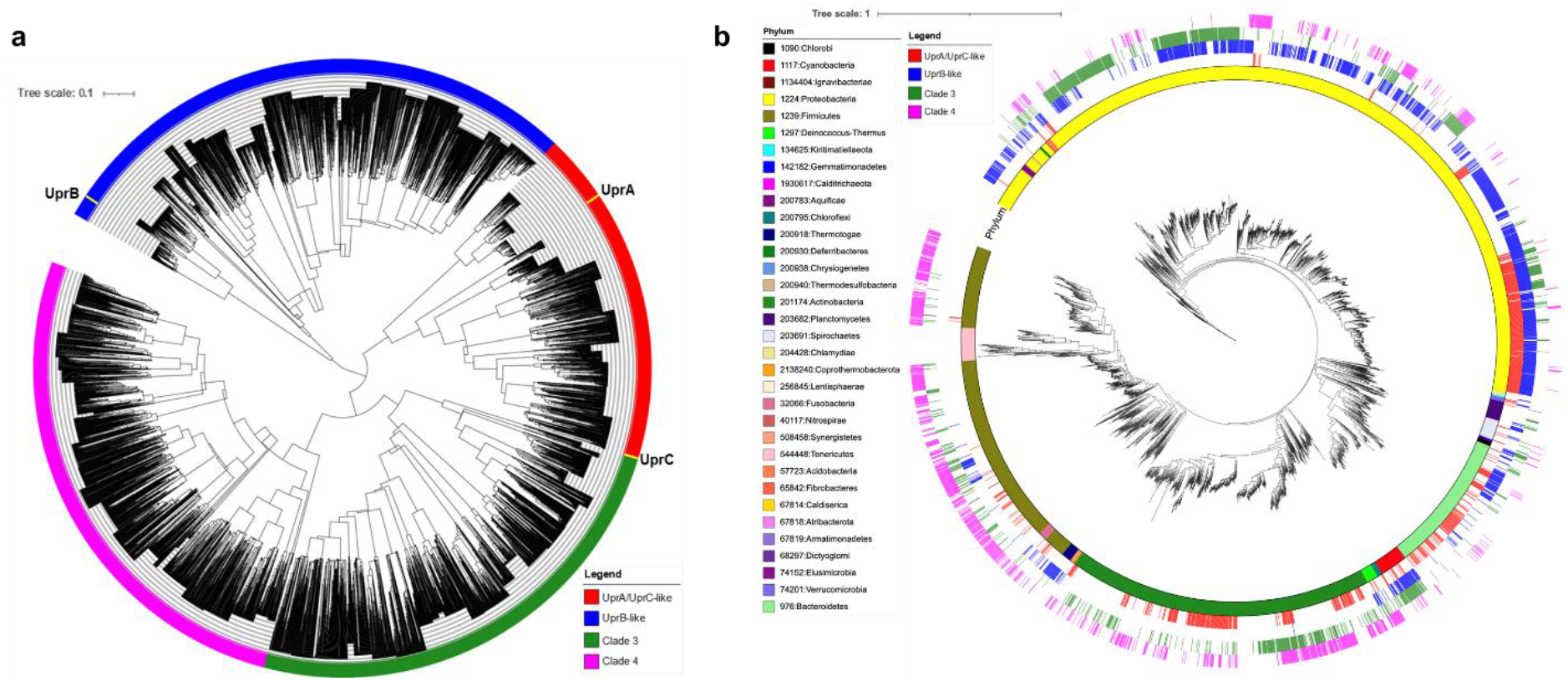
Upr-like enzymes are widespread amongst bacteria. (**a**) Phylogenetic tree of Upr-like protein sequences. The tree was constructed from the 4860 Upr-like sequences found in the 3992 diverse bacterial genomes mentioned in Figure 6. Upr-like enzymes are separated into 4 clades: UprA/UprC-like (red), UprB-like (blue), Clade 3 (green), and Clade 4 (magenta). (**b**) Phylogenetic tree showing distribution of the four clades of Upr-like proteins in the diverse set of 3992 bacterial taxa mentioned in Figure 6.

**Supplementary Table S1.**
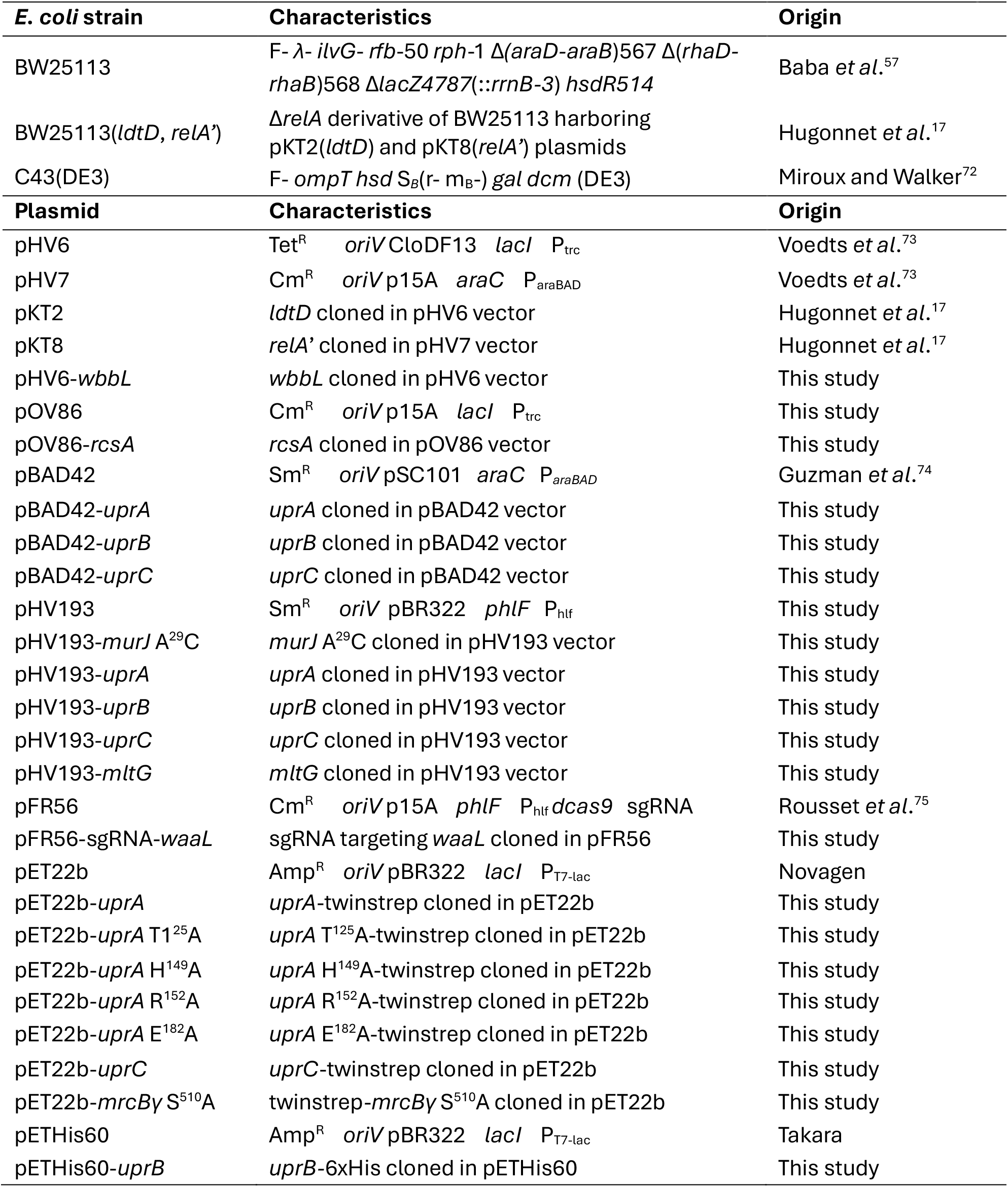
Characteristics and origin of strains and plasmids.

